# On the bacterial ancestry of mitochondria: new insights with triangulated approaches

**DOI:** 10.1101/2022.05.15.491939

**Authors:** Michelle Degli Esposti, Otto Geiger, Alejandro Sanchez-Flores, Mauro Degli Esposti

## Abstract

We breathe at the molecular level when mitochondria in our cells consume oxygen to extract energy from nutrients. Mitochondria are characteristic cellular organelles that derive from aerobic bacteria similar to some of those thriving in the oceans nowadays. These organelles carry out most metabolic pathways in eukaryotic cells. The precise bacterial origin of mitochondria and, consequently, the metabolic ancestry of our cells remains controversial - despite the vast genomic information that is now available. Here we triangulate across multiple phylogenomic and molecular approaches to pinpoint the most likely living relatives of the ancestral bacteria from which mitochondria originated.

## INTRODUCTION

Unveling thre origins of mitochondria continues to challenge science. While there is broad consensus that mitochondria first evolved around 1700 million years ago, the critical question of from which bacteria they originated remains unanswered [1–11]. Previous research has primarily relied on phylogenetic inference to identify the possible bacterial relatives and ancestors of mitochondria, hereafter called protomitochondria. However, this approach has produced inconsistent and varying results depending on the phylogenetic approach, taxonomic sampling, and corrections used to reduce artefacts [1–8]. In fact, almost all major lineages of alphaproteobacteria have been proposed as protomitochondria relatives. The fragility and inconclusiveness of available evidence suggests that phylogenetic trees may not be sufficient for identifying the extant bacteria that are closest to protomitochondria. This is likely due to the vast amount of time passed since the original symbiotic event, which has diluted and dispersed the phylogenetic signal of contemporary bacterial proteins with respect to their mitochondrial homologs [2,6]. Moreover, the debate whether the bacterial ancestor of mitochondria was an obligate or facultative aerobe (see [10] for a recent review) further complicates the evaluation of the metabolic ancestry of protomitochondria. Thus, new and robust evidence is needed to unveil the origins of mitochondria [6,10,11].

To provide such evidence, here we triangulate across multiple phylogenomic and molecular approaches with different sources of biases [12], covering aerobic and anaerobic metabolic traits shared by bacteria and mitochondria [6–11] (Table 1). The guiding principle of this strategy is that the creation of the first eukaryotic cell involved genomic transmission of metabolic traits from a bacterium that could have surviving descendants today. Although the transmission has been a rare, if not singular event [1,3,6,10], it might have left vestigial traces in the genome of some of those descendants – similar to ‘missing link’ features found in other major transitions in evolution. An example of evolutionary traces of this kind is the synteny of two genes of cytochrome *c* oxidase (COX or complex IV) [11,13], the mitochondrial enzyme which ultimately consumes oxygen in our cells (Fig. 1 and Supplementary Fig. S1). Seven genes for COX subunits and accessory proteins form a conserved genomic cluster (operon) that is characteristic of alphaproteobacteria [11]. Four of these genes are encoded in mitochondrial DNA (mtDNA) of early branching unicellular eukaryotes [11,13–15], while two others are present in mitochondrial complex IV of the protist *Tetrahymena* [16] (Fig. 1). Moreover, the gene for the assembly protein Cox11 (Cox11_CtaG) always precedes the gene for the COX3 subunit (Figure 1), forming a collinearity that is conserved in the mtDNA of some protists (Supplementary Fig. S1). The Cox11-COX3 synteny can thus be considered a genomic relic of the aerobic ancestry of protomitochondria, providing a selection criterion for putative bacterial relatives of mitochondria [11]. Its absence in the genome of most bacteria (Fig. 1), including the Rickettsiales often considered as relatives of mitochondria [4,7,13], would exclude such prokaryotes from the ancestry of protomitochondria. Here we present diverse new approaches that confirm this exclusion and indicate that the ancestor of protomitochondria was likely related to extant alphaproteobacteria never considered before for the evolution of mitochondria.

**Table 1.** The main approaches of the triangulation strategy for defining the bacterial ancestry of mitochondria.

| Approach and criterion | Major sources of bias | Direction and influence of biases |  |
| --- | --- | --- | --- |
| No difference in cumulative scores of aerobic traits | ● favors larger genomes<br>● multiple genes for same trait inflate scores | ↑ | scores *increase* with genome size approaching aerobic mitochondria |
| presence of *spt* and *kynU* genes for ceramide and kynurenine | ● gene loss disfavors presence<br>● lack of biochemical evidence for trait disfavors presence | ↓ | absence of genes may derive from differential loss subsequent to protomitochondria evolution |
| presence of anaerobic traits for quinones and ferredoxin oxido-reductases | ● favors facultative aerobes<br>● increases with gene duplication | ↑ | genes *increase* in number by ecological adaptation to anoxia as in anaerobic eukaryotes |
| very few INDELs in respiratory proteins | ● genome streamlining reduces the number of INsertions | ↓ | INDELs *decrease* by convergent evolution with mitochondria |

**Figure 1.**
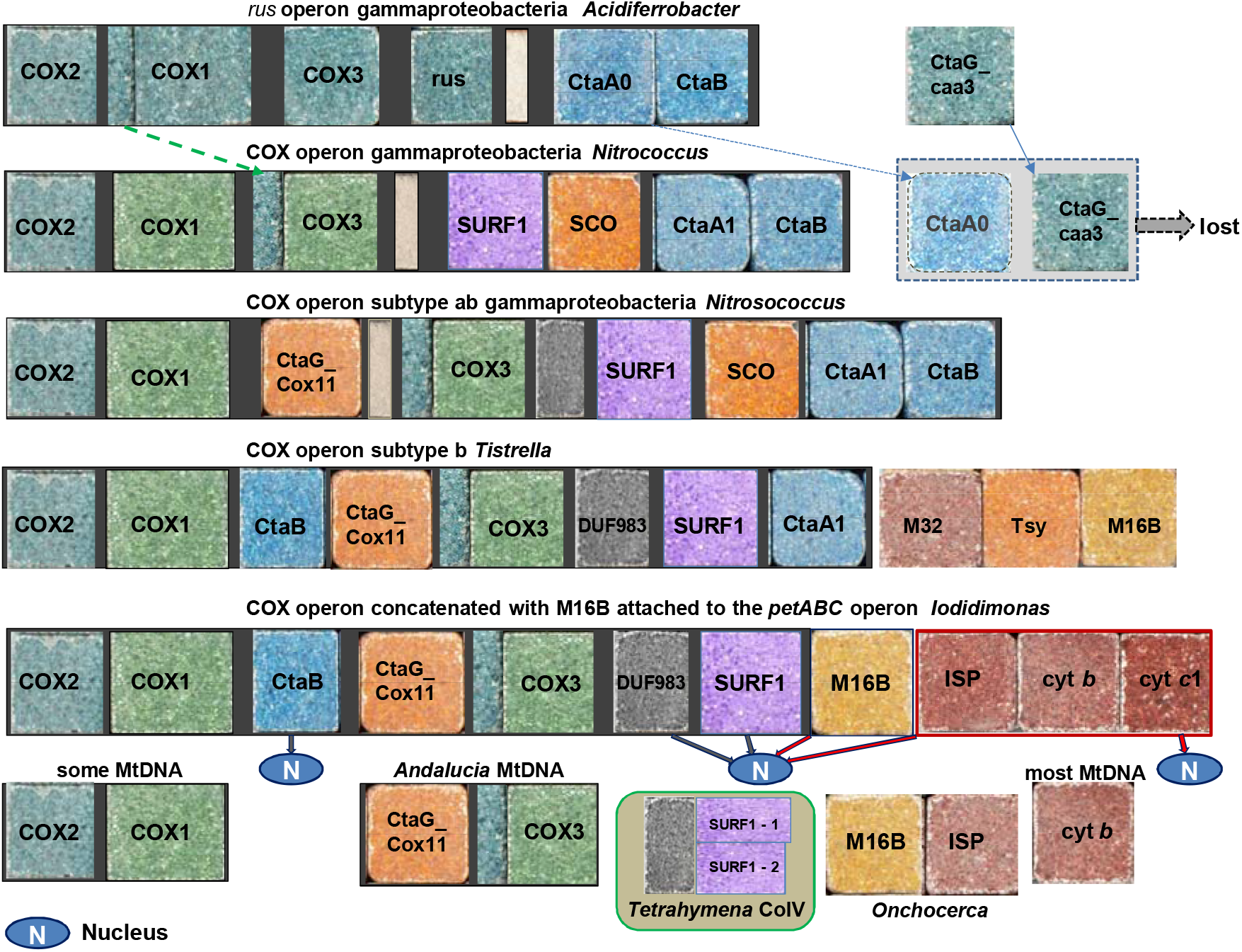
Evolution of the cytochrome oxidase gene cluster from proteobacteria to mitochondria. The figure illustrates the gene clusters (operons) of cytochrome *c* oxidase rendered with Roman mosaic tiles to indicate the mosaic nature of the proteins encoded by the clustered genes [11,13,30,43] and of the whole mitochondrial proteome [3,62]. The *rus* operon of iron-oxidizing bacteria such as *Acidiferrobacter* (first row in the illustration) is the archetype of COX operons of A1 type [43] and includes an ancestral form of both Heme A synthase (CtaA0) and transmembrane CtaG (CtaG_caa3). These genes are retained in the genome of nitrifying *Nitrococcus* [43] (second row in the illustration), but have been subsequently lost along COX evolution, substituted by CtaA1 and CtaG_Cox11, as shown in the operon of *Nitrosococcus* (third row in the illustration). The transition from the *rus* operon to the COX operon of *Nitrococcus* included the increase in the molecular size of ancestral COX3, which originally had five transmembrane helices (TM) and then acquired two additional TM at its N terminus, thereby forming the 7 TM architecture which is seen in both alphaproteobacterial and mitochondrial structures [16,18,20,63]. Our sequence analyses suggest that the two extra TM present at the N terminus of ancestral COX1 [43] might have constituted the source for the additional two TM fused to the COX3 core, as indicated by the dashed green arrow on the top of the illustration. The COX operon of *Tistrella* [13] is intermediate between that of gammaproteobacteria such as *Nitrosococcus* and that typical of other alphaproteobacteria [11]. The gene for M16.06 group of M16 zinc peptidases [22] is often associated to the end of alphaproteobacterial COX operons, generally together with the gene for threonine synthase (Tsy) after the SURF1 gene. DUF983, a domain of unknown function protein with two closely spaced TM at its centre [11], generally precedes the SURF1 gene in the same operon. Intriguingly, potential homologs of bacterial DUF983 proteins and two SURF1 isoforms are part of *Tetrahymena* Complex IV [16]. This is represented by the central box in the last row of the illustration, which summarizes the genomic transition between alphaproteobacteria, with the rare concatenation of the COX operon with M16B gene plus the *petABC* operon of the *bc*_1_ complex of *Iodidimonas* [64], and eukaryotes. The last row includes a protein from the filarial *Onchocerca* (Supplementary Fig. S2) containing the collinear fusion of M16B with ISP encoded in the nuclear DNA of eukaryotes (N symbol). Similar fused proteins have been found in two other nematodes.

## RESULTS and DISCUSSION

### 1. Unveiling a new synteny in complex III genes

We searched for other genomic traces with equivalent discriminating power as the Cox11-COX3 synteny, focusing on the possible genomic association of two proteins that are structurally and functionally interconnected in complex III: the Mitochondrial Processing Protease (MPP) and the Rieske Iron Sulfur Protein (ISP, Fig. 1). Mitochondrial complex III (ubiquinol:cytochrome *c* reductase) derives from the bacterial cytochrome *bc*_1_ complex, which is encoded by the *petABC* operon now represented by the cytochrome *b* gene of mtDNA (Fig. 1, cf. [13,17]). In plants and protists, two MPP proteins form a large domain of complex III structure, not present in bacterial *bc*_1_ [16,18,19]. In animal mitochondria, MPP derivatives called Core Proteins (CP) have the same structural organization [17,20], while the functional MPP heterodimer is a separate soluble enzyme [18]. We have confirmed two genes for MPP proteins but hardly any for CP proteins in the genomes of Rhodophyta, Discoba and other early branching eukaryotes (Supplementary Table S1). Hence, it is highly likely that MPP proteins were constitutive components of the first mitochondrial complex III, as seen in *Tetrahymena* [16]. Notably, CP retain the function of processing the pre-sequence of ISP [17,20], thereby underlying the intimate connection between MPP and ISP. Following the finding of a filarial protein corresponding to the fusion of MPP with ISP (Fig. 1), we systematically searched the genomes of currently available bacteria for the contiguity of genes encoding the bacterial homologs of MPP and ISP. The homolog and likely precursor of MPP is a zinc peptidase belonging to a specific group of the M16B subfamily [21,22]. We found its gene adjacent to that of ISP in two species of *Iodidimonas, I.muriae* (Fig. 1) and *I. gelatinilytica* (Supplementary Table S2), as well as in related alpha proteobacteria Q-1 (Supplementary Fig. S2). *Iodidimonas* is part of the SERIK group, including the orders of Sneathiellales, Emcibacterales, Rhodothalassiales, Iodidimonadales and Kordiimonadales [2]. The few genomes in which MP16B and ISP are separated by one to five other genes belong to SERIK taxa too (Supplementary Fig. S2).

### 2. Distribution of aerobic traits in mitochondria and alpha proteobacterial lineages

The rare M16B-ISP synteny (Fig. 1) would represent a novel trace of the metabolic ancestry of protomitochondria or may derive from some unusual genomic streamlining. To discriminate between these possibilities and build a thorough strategy of triangulation [12], we followed different approaches and considered their potential biases as well as the expected direction of such biases (Table 1). In the first approach, we systematically analysed 18 traits of mitochondrial aerobic metabolism which are variably distributed among the principal lineages of alphaproteobacteria (Box1). The traits include proteins encoded in the characteristic COX operon, to which M16B is frequently associated (Fig. 1), as well as other proteins that participate in the assembly of complex IV (SCO, CtaA/Cox15 [23]), complex III (Cbp3 chaperone [23]) and their substrate cytochrome *c* (heme lyase CcmF). Among traits not evaluated before, we considered a bacterial zinc-finger protein that is the likely ancestor of subunit Vb of complex IV [24] and a cysteine signature in a conserved C-terminal region of COX1 [25] (Box1). The latter was considered for its common presence in alphaproteobacteria with reduced genomes including Rickettiales and early branching eukaryotes (Supplementary Table S2). We also evaluated the distribution of genes for M16A peptidases, which have multiple homologs in eukaryotes [26] (Box1; see Methods for more detail), but have not been considered before in relation to eukaryogenesis [1–8,13]. Figure 2 presents the lineage-specific distribution of the selected aerobic traits in quantitative terms, showing an apparent peak around Iodidimonadales. This peak depends only partially on the premium given to the presence of the M16B-ISP synteny and is not influenced by the major bias of our first approach, namely genome size (Table 1). Indeed, Iodidimonadales have genomes smaller than the average genome of alphaproteobacteria. In any case, the cumulative aerobic traits score for Iodidimonadales does not significantly differ from aerobic mitochondria (Fig. 2 and Supplementary Table S2B). Sneathiellales, Kordiimonadales, Sphingomonadales and Caulobacterales also show no statistically significant difference in the distribution of aerobic traits in comparison with mitochondria (Fig. 2 and Supplementary Table S2B). In contrast, the lineages of MarineProteo1, MarineAlpha, Rickettsiales, Holosporales, Pelagibacterales and Rhodobacterales have significantly lower cumulative scores than aerobic mitochondria (Fig. 2), suggesting their exclusion from the aerobic ancestry of protomitochondria. Conversely, the distribution of aerobic traits in other lineages partially overlaps that of mitochondria (Fig. 2). It is thus difficult to exclude such lineages from potential ancestors of protomitochondria based on statistical distributions alone. Different approaches are needed to overcome specific sources of bias and further filter alphaproteobacteria lineages for identifying the most likely bacterial ancestor of protomitochondria (Table 1).

**Figure 2.**
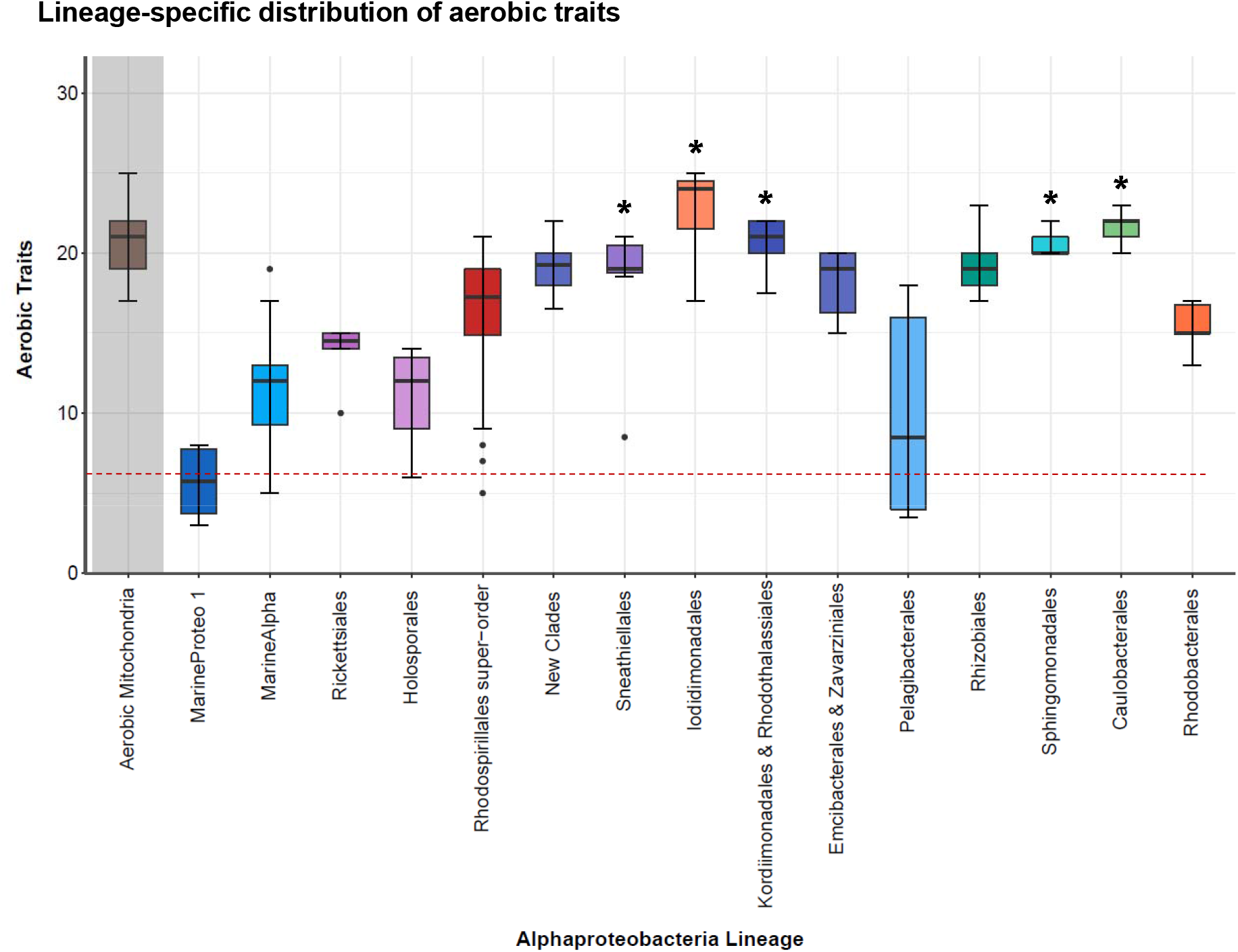
Distribution of aerobic traits scores along alphaproteobacterial lineages and mitochondria. The figure presents the lineage-dependent plot of the individual cumulative scores for the 18 aerobic traits considered here (Box1) in their branching order along the x axis (except for the mitochondria on the left). Asterisks indicate distributions of aerobic traits that are not significantly different from that of aerobic mitochondria (p>0.1 with the 99% Confidence T Test, Supplementary Table S2B). Note that the lineage of MarineProteo1 lacks the A1 type COX1 and COX2 proteins that are shared by alphaproteobacteria and mitochondria [11]. The dashed red line at the bottom of the plot indicates the median value of cumulative scores for the same aerobic traits obtained in the genome of gammaproteobacteria such as *Nitrococcus* and *Nitrosococcus*, cf. Fig. 1.

#### Box1 List of the aerobic traits considered for the analysis in Fig. 2 - see also Methods section.

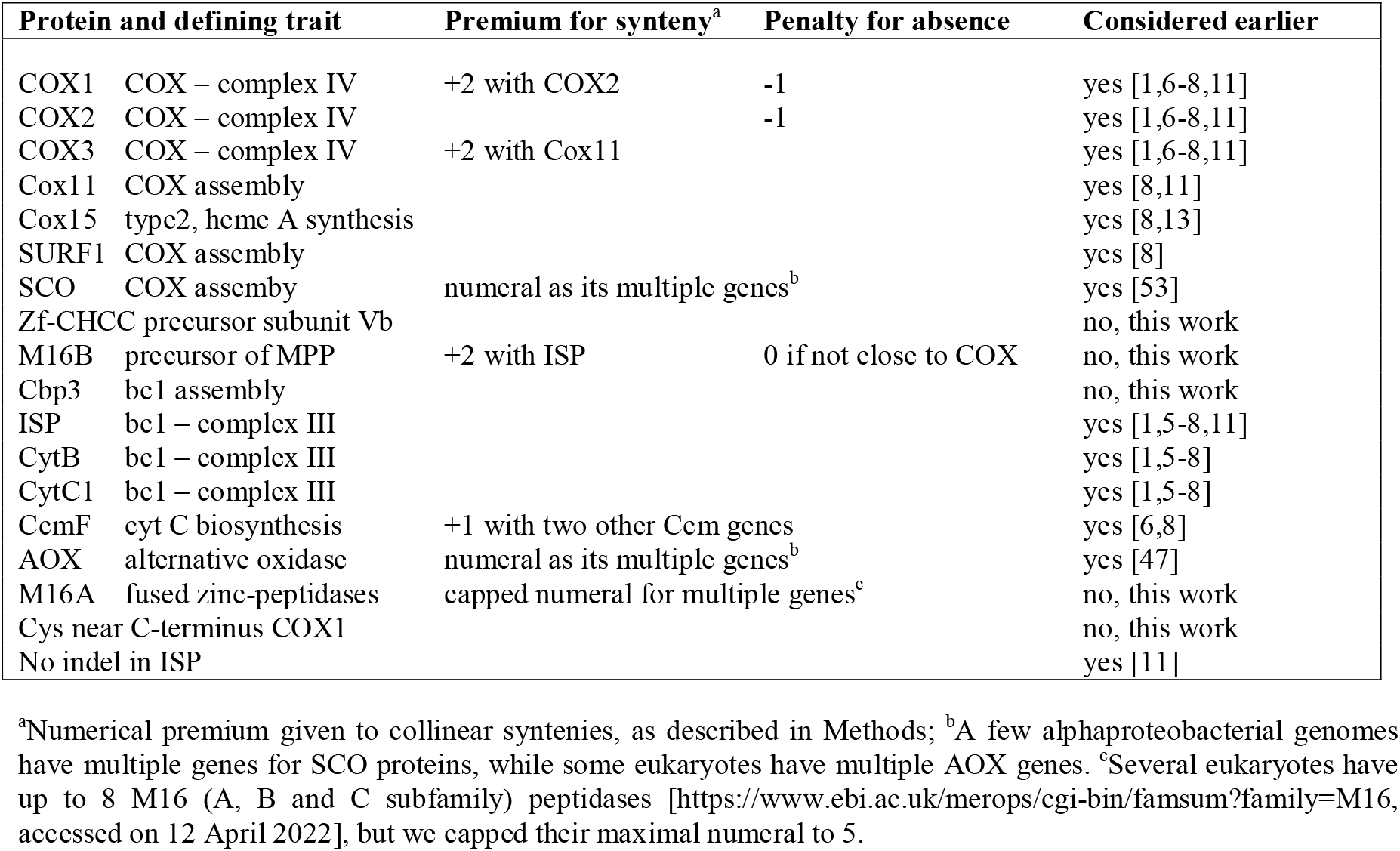

### 3. Distribution of genes for ceramide and kynurenine biosynthesis in alphaproteobacteria

Our next approach was based on a completely different metabolic pathway which has hardly been considered before in regard to eukaryogenesis: the biosynthesis of ceramide-based lipids, sphingolipids. Sphingolipids constitute a vast class of membrane lipids that are ubiquitous in eukaryotes, but scarcely present in bacteria [27,28]. To produce the precursor of ceramide, bacteria often require a four-genes operon ending with the gene for an α-oxoamine synthase catalyzing the key step of ceramide biosynthesis: serine palmitoyl-transferase (SPT) (Fig. 3A, cf. [28,29]). We found the *spt* gene encoding SPT in Iodidimonadales and other members of the SERIK group, as well as in some Rhodospirillales and a few Rhizobiales (Figs. 3 and 4, Supplementary Table S3). Our phylogenetic analysis indicates that *Odyssella* sp. NEW MAG-112 may have the earliest enzyme for ceramide biosynthesis of all alphaproteobacteria (Fig. 3). Alphaproteobacterial SPT is clearly the ancestor of both isoforms of eukaryotic SPT (Fig. 3B), as well as of nitrifying taxa such as *Nitrococcus* (Fig. 3C). Intriguingly, such nitrifying bacteria possess intra-cytoplasmic membranes resembling mitochondrial cristae, as in alphaproteobacterial methanotrophs (Methylocystaceae [30,31]) that also have SPT (Fig. 3C). Ceramide is well known to modulate the curvature and shape of lipid bilayers [32] and may therefore be crucial for the formation of bacterial intracytoplasmic membranes. The genomic distribution of *spt* and its partner genes shows a maximum within the SERIK group, besides the expected high frequency in Sphingomonadales (Fig. 4). The limited presence of the same genes in different lineages probably derives from events of lateral gene transfer (LGT), since they are present only in a few taxa that have SPT proteins clustering with those of other alphaproteobacteria, as in the case of *Caulobacter* (Fig. 3C). We also found that SPT distribution often matches that of the kynureninase gene *kynU* (Supplementary Table S3), another pyridoxal 5’-phosphate-dependent enzyme defining the kynurenine pathway for NAD(P)^+^ biosynthesis in bacteria [33]. An equivalent pathway is required for *de novo* synthesis of rhodoquinone (RQ) in nematodes and other eukaryotes, which use this quinone in their adaptation to low levels of oxygen [34] (see below). The *kynU* gene is generally associated with *kynA* encoding tryptophan dioxygenase, the upstream enzyme of the kynurenine pathway [33,34]. This genomic association is present in several taxa of the SERIK group but absent in Caulobacterales (Supplementary Table S3). Hence, the combination SPT-kynureninase produces a stringent criterion for discriminating alphaproteobacterial lineages from the ancestry of protomitochondria, which must have had both traits that are now present in a variety of eukaryotes.

**Figure 3.**
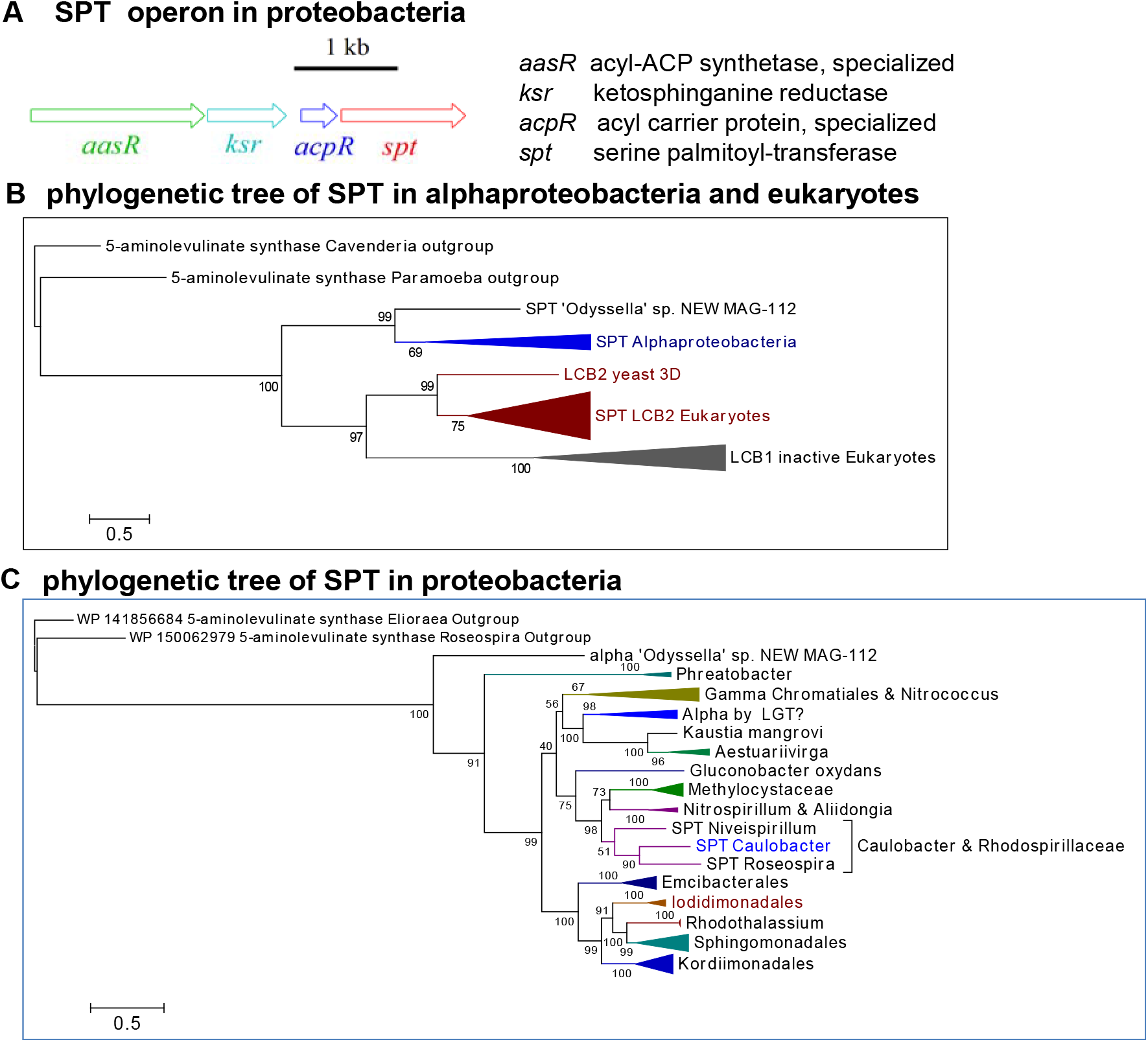
Genomics and phylogenetic trees of bacterial and eukaryotic SPT. A. Representation of the four-genes operon comprising SPT found in proteobacteria, for example *Caulobacter* [27,29]. B. Phylogenetic ML tree of SPT from alphaproteobacteria and various Eukaryotes, which have two isoforms [28]: the catalytic LCB2 and the inactive LCB1. Paralog proteins of 5-aminolevulinate synthase are used as outgroup providing the root of the tree. The alpha MAG originally named *Odyssella* sp. NEW MAG-112 (GCA_016792765.1) is not a member of the Holosporales, but of the order o_Bin65 according to GTDB taxonomy [58]. It is included here in the lineage of Emcibacterales & Zavarziniales for similar INDELs profiles (Supplementary Tables S2 and S4), and therefore labeled as ‘Odyssella’. C. ML tree of proteobacterial SPT proteins, including nitrifying gammaproteobacteria such as *Nitrococcus*. Note the clustering of *Caulobacter* SPT with homologs of Rhodospirillaceae, suggesting it may derive from LGT, considering also that other Caulobacterales genera do not have the trait - except for a few *Phenylobacterium* taxa (Supplementary Table S3). The SPT of the betaproteobacterium *Nitrosomonas* [29] clusters with those of gammaproteobacteria, as found in separate ML trees. Numbers indicate the strength of the nodes in percentage values of ultrafast bootstraps.

**Figure 4.**
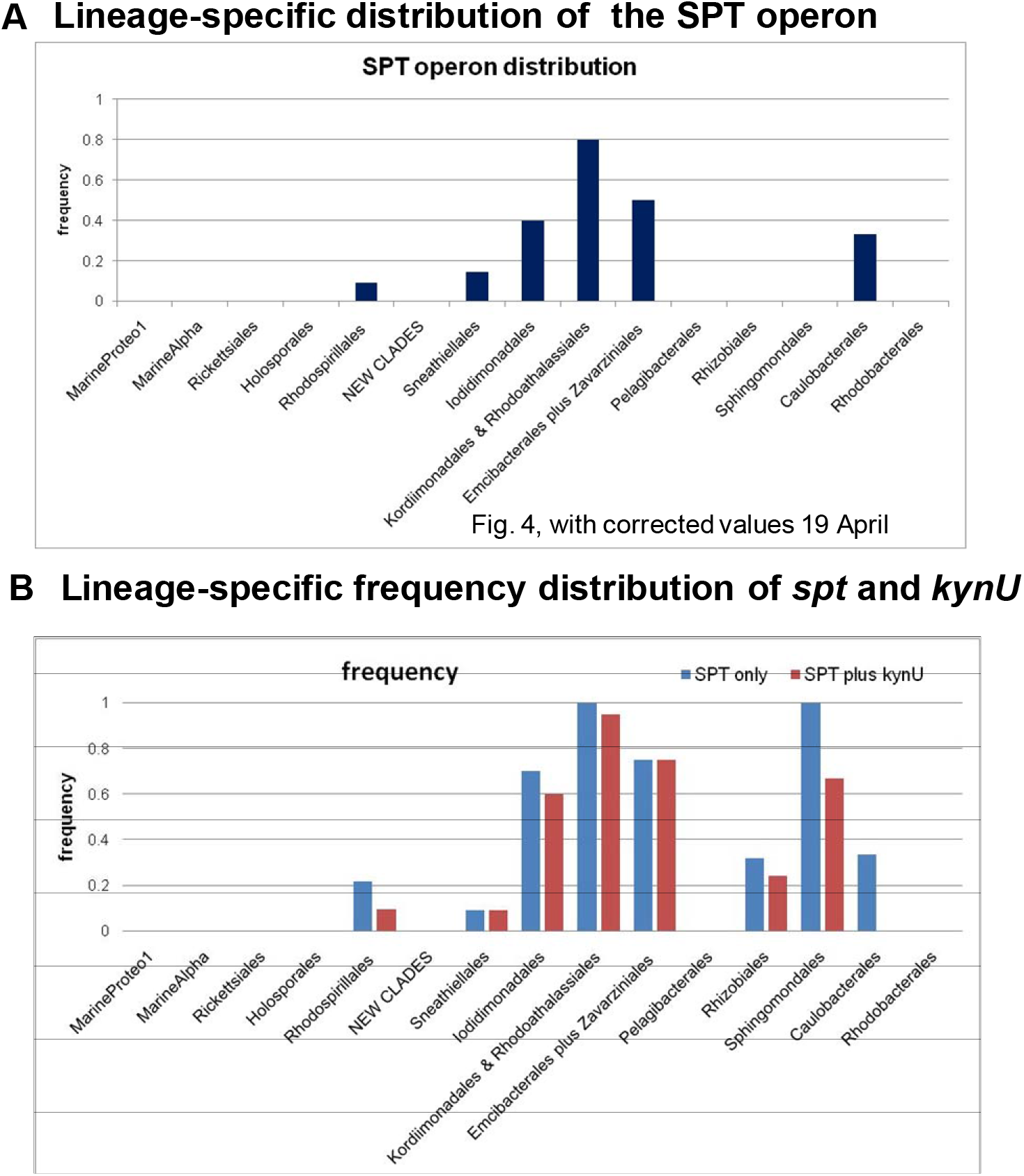
Distribution of the SPT operon and *spt* plus *kynU* genes along alphaproteobacterial lineages. A. Distribution of the four-genes *spt* operon as determined by detailed genomic searches (Supplementary Table S3). Note that Sphingomonadales, which are well known to produce ceramide lipids [27,28], are not represented because they have different variants of the operon with at least one of its genes displaced in other genomic regions [27,29]. B. Frequency distribution of the combination of the *spt* and *kynU* genes, irrespective of their surrounding genes, along the alphaproteobacterial lineages. Note that the majority of the eukaryotic taxa used for the analysis in Fig. 2 do have SPT proteins (Fig. 3B), while the distribution of the *kynU* gene is scattered [34].

### 4. Analysis of anaerobic traits along alphaproteobacteria lineages

The previous analysis of *kynU* distribution has suggested that *de novo* synthesis of RQ may occur also in alphaproteobacteria. This would be an important novelty relevant to the bacterial ancestry of protomitochondria, because RQ is one rare trait of anaerobic metabolism that is shared by mitochondria and alphaproteobacteria [34–36] (see Materials and Methods for more details). Members of the Azospirillaceae family that have *kynU* and related genes of the kynurenine pathway (Supplementary Table S3) also have two or more genes encoding different forms of the UbiA protein catalysing a critical step in ubiquinone (Q) biosynthesis (Figure 5). This is an infrequent occurrence that echoes a distinctive feature of *de novo* RQ biosynthesis in *C.elegans* and other metazoans: the presence of a second UbiA gene for a protein that specifically transfers the isoprenoid tail to a ring precursor derived from the kynurenine pathway [34]. Therefore, the presence of multiple UbiA genes in the genome of alphaproteobacteria that likely have the kynurenine pathway (boxed in Figure 5) suggests that such bacteria may synthesize RQ by a *de novo* system equivalent to that of *C.elegans*. We additionally discovered that the genomes of several such alphaproteobacteria also possess the *ubiTUV* genes for the anaerobic biosynthesis of Q [37,38] (Fig. 5). These are: *Nitrospirillum amazonense, Arenibaculum*, most members of the *A.lipoferum* clade, *Rhodothalassium* spp. and four MAGs including *Odyssella* sp. NEW MAG-112, which has an early branching SPT (Fig. 3B). Notably, the proteins encoded by *ubiU* and *ubiV* bind a 4Fe4S cluster promoting the oxygen-independent hydroxylation of the Q ring under anaerobic conditions [37]. Under normal oxygen conditions, this hydroxylation is catalyzed by flavin hydroxylases such as UbiH [38]. Intriguingly, we have found paralogs of UbiU in Chlorophyta and other protists, often in fused proteins containing a similar, UbiV-like domain (see Materials and Methods for more details). It is thus possible that the *ubiU-ubiV* synteny has been transmitted by an alphaproteobacterial progenitor to primordial eukaryotes, thereby rendering these genes powerful new markers for the anaerobic metabolism shared by facultatively aerobic bacteria and eukaryotes [2,35,37,38].

**Figure 5.**
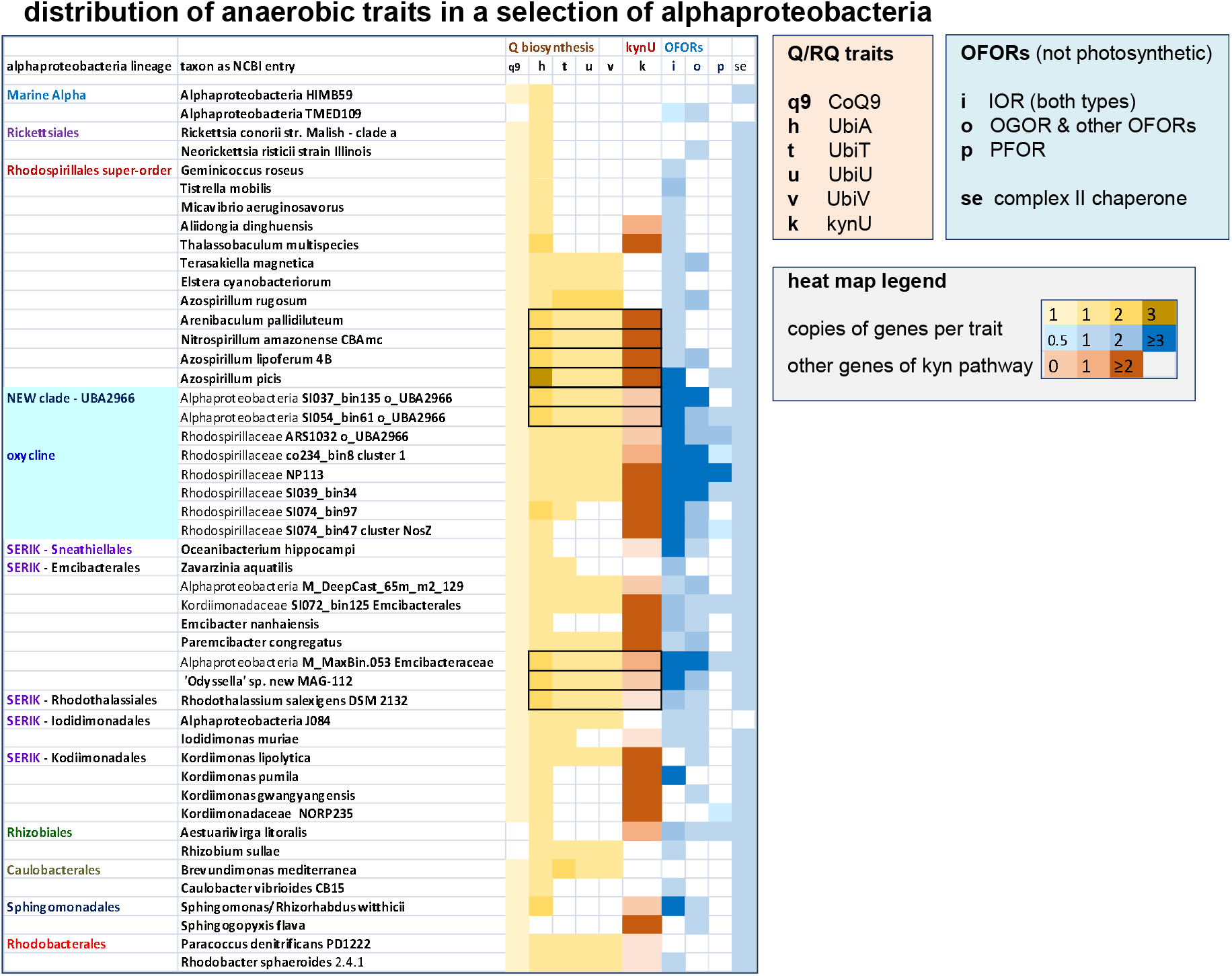
Distribution of the anaerobic traits in a selection of alphaproteobacteria. At least two representatives for each of the lineages of alphaproteobacteria considered here (cf. Fig. 2) are presented in a compacted phylogenetic sequence from top to bottom. Several taxa were considered for Rhodospirillales and SERIK group lineages, as well as the lineage of New clades [2], because they presented the highest concentration of OFORs traits (Supplementary Fig. S3) as well as those for the anaerobic production of Q [2,37,38]. Traits boxed in black squares indicate the potential for *de novo* biosynthesis of RQ (see text). The abbreviations for the various traits are described in the legend boxes on the right. The traits of CoQ9 (q9) on the far left and of the chaperone of complex II (se) on the far right are used as reference common genes shared with mitochondria.

Next, we considered the distribution of 2-oxoacid ferredoxin oxidoreductases (OFORs) as established traits for anaerobic metabolism [7,35]. Five different types of OFORs [39] are found in the genomes of alphaproteobacteria, the long indolepyruvate:ferredoxin oxidoreductase (IOR) being the most common (Fig. 5 and Supplementary Fig. S3A). The cumulative distribution of OFORs genes is uneven along the various lineages, with maximal concentration in marine New clades [2] (Fig. 5 and Supplementary Fig. S3). Eukaryotes adapted to anoxic conditions predominantly have the pyruvate:ferredoxin oxidoreductase (PFOR) and the oxoglutarate:ferredoxin oxidoreductase (OGOR) in hydrogenosomes or other mitochondria-related organelles [35]. Interestingly, the genes for these proteins are concentrated in the phylogenetic space spanning the New clades and the SERIK group [2], which also show the highest scores for aerobic traits (Fig. 5 and Supplementary Fig. S3B). Hence, the distribution of the metabolic traits considered here is not random among extant alphaproteobacteria, converging on a central phylogenetic region that may have the highest probability for protomitochondria ancestry. To investigate this possibility, we used a completely different approach (Table 1).

### 5. INDEL analysis of respiratory proteins

The convergence of the results obtained with the approaches used so far (Figs. 2–5) appears to strengthen the pivotal position of the SERIK group [2] for the ancestry of protomitochondria. However, it cannot exclude different alphaproteobacteria such as *Nitrospirillum*, which may have the capacity of *de novo* biosynthesis of RQ and possess other anaerobic traits (Fig. 5). Moreover, no member of the SERIK group has been previously considered as a possible ancestor of protomitochondria [1–11,13].To strengthen our novel findings, we developed a different triangulation approach based on conserved complex INDELs (INsertions or DELetions [40–42]), which follows principles of molecular rather than phylogenomic analysis. The identification of conserved INDELs emerged from detailed alignments of the catalytic subunits of both complex III and IV (Supplementary Table S4). Three of such INDEls are present in COX3 and have evolutionary value for they are shared with *Nitrococcus* COX3 (Supplementary Fig. S4), which occupies an intermediate position in the evolution of the COX operon from its ancestral form in iron-oxidizing bacteria of the same gammaproteobacteria class (Fig. 1 – cf. [43]). These INDELs are retained in COX3 sequences of alphaproteobacteria such as Caulobacterales, while they disappear in the proteins of *Magnetospirillum* and related taxa, which have a simplified COX operon lacking both CtaB and SURF1 [11]. Such alphaproteobacteria, therefore, are unlikely to be the source of the various COX proteins of mitochondria (Box1). The absence of INDELs can be considered the ancestral state of mitochondrial COX3 and occurs also in *Iodidimonas*, but not Azospirillaceae (Supplementary Fig. 4). Given the phylogenetic limits of complex INDELs shared by multiple bacterial lineages [41,42], we expanded our analysis by including well defined INDELs in the bacterial sequences of the NuoL and NuoD subunits of complex I (Figure 6). Mitochondrial sequences have gaps corresponding to the bacterial INDELs examined; therefore, in principle, the lower the cumulative count of INDELs, the higher the possibility of being related to protomitochondria. However, this principle is tempered by the caveat of genome streamlining, which reduces the number of protein insertions due to parsimony, thereby producing a different directional bias in comparison to mitochondria (Table 1). For example, Pelagibacterales have an extremely streamlined genome [44] and show a significant reduction in the cumulative count of INDELs (Fig. 6) with respect to phylogenetically close lineages such as Rhizobiales [2,7]. The comparable low counts in other taxa with reduced genome [1,8], for example Rickettsiales (Fig. 6), has little relevance to the ancestry of mitochondria, because we have already excluded such taxa using previous approaches (Figs. 2–5). Considering the wide variance of the cumulative data (Fig. 6, grey histograms) and the potential randomness of INDELs [42], a stringent discriminatory threshold could be set at 3, above the standard deviation of Pelagibacterales. This threshold excludes Kordiimonadales, Emcibacterales, Rhizobiales, Sphingomonadales and, again, Caulobacterales plus Rhodobacterales (Fig. 6).

**Figure 6.**
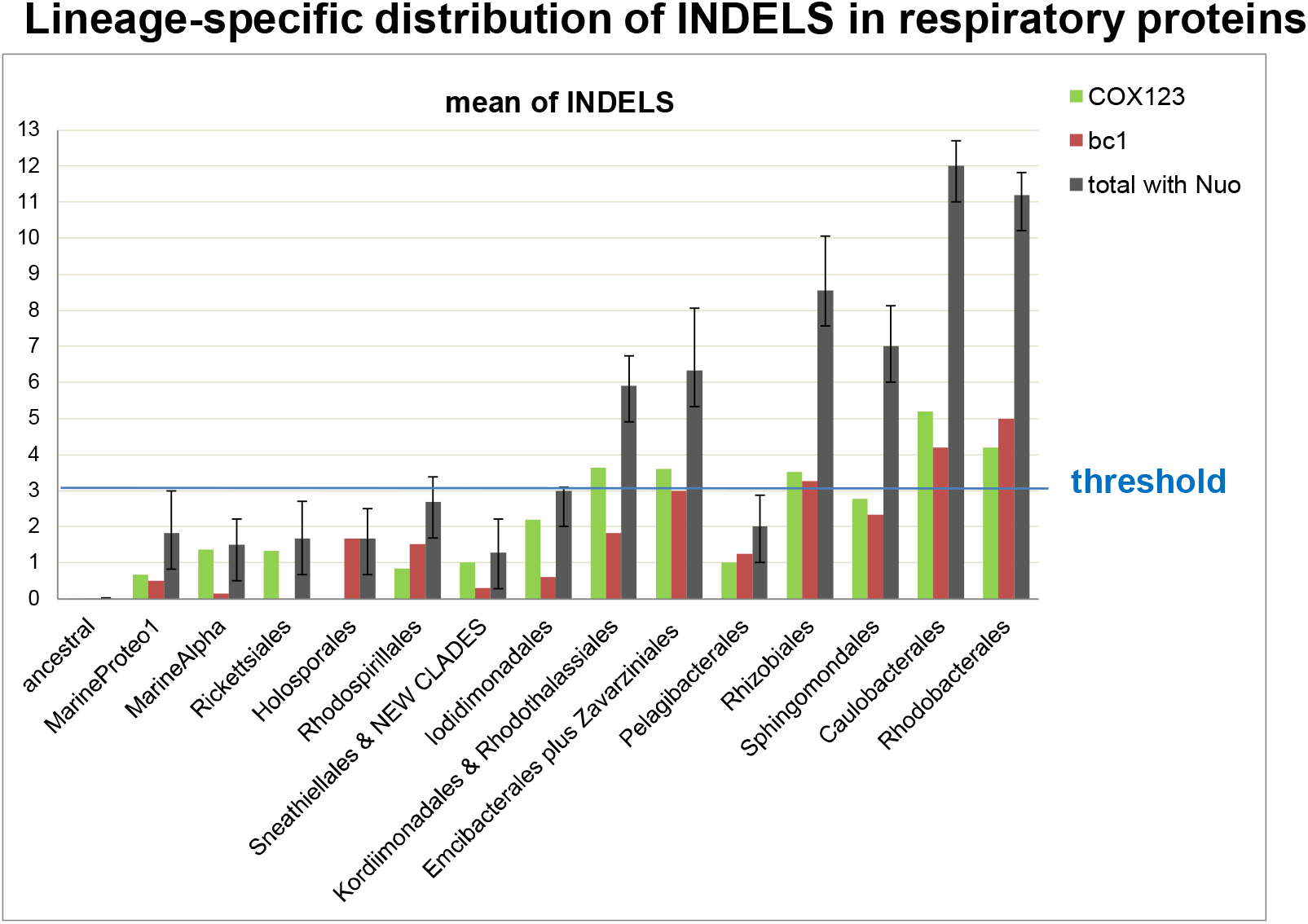
Distribution of conserved INDELs in 8 respiratory proteins along alphaproteobacterial lineages. The plot shows the mean plus Standard Deviation (SD, represented by the error bars) of the conserved INDELs found in the catalytic subunits of complex III and IV, plus two subunits of complex I (Supplementary Table S4) along alphaproteobacterial lineages. At difference with previous plots, the lineage of New clades (labelled as NEW CLADES) is amalgamated to that of Sneathiellales because there is no statistically significant difference in the INDELs distribution of the separate lineages. The blue line indicates a threshold just above the upper SD of Pelagibacterales, as discussed in the text. Note that the mitochondrial homolog proteins of the eukaryotic taxa in Fig. 2 do not have these bacterial INDELs (cf. Supplementary Fig. S4A) and are annotated as ancestral on the left.

### 6. Different approaches indicate that *Iodidimonas* spp. may be close to protomitochondria

Table 2 summarizes the results obtained with the different approaches evaluated here for screening bacteria that may be close to protomitochondria (Table 1), plus a criterion based upon collinearity of ribosomal genes [13]. *Tistrella*, previously considered a possible relative to protomitochondria [11,13], shows this collinearity and no conserved INDELs (Supplementary Table S4), but does not have SPT nor many anaerobic traits (Table 2). Together, our results indicate that very few alphaproteobacteria would be positively selected by more than three triangulated approaches (Table 2). *Iodidimonas* spp. are superior to other possible candidates because they are consistently selected by widely different approaches (Tables 1 and 2, cf. [12,45]). We thus hypothesize that *Iodidimonas* spp. may be descendant of the ancestral bacteria that originated protomitochondria. Indeed, they are related to metagenome assembled genomes (MAGs) uncovered in a hydrothermal environment recapitulating oxygen evolution in proterozoic earth [46] (Supplementary Fig. S4B).

**Table 2.**
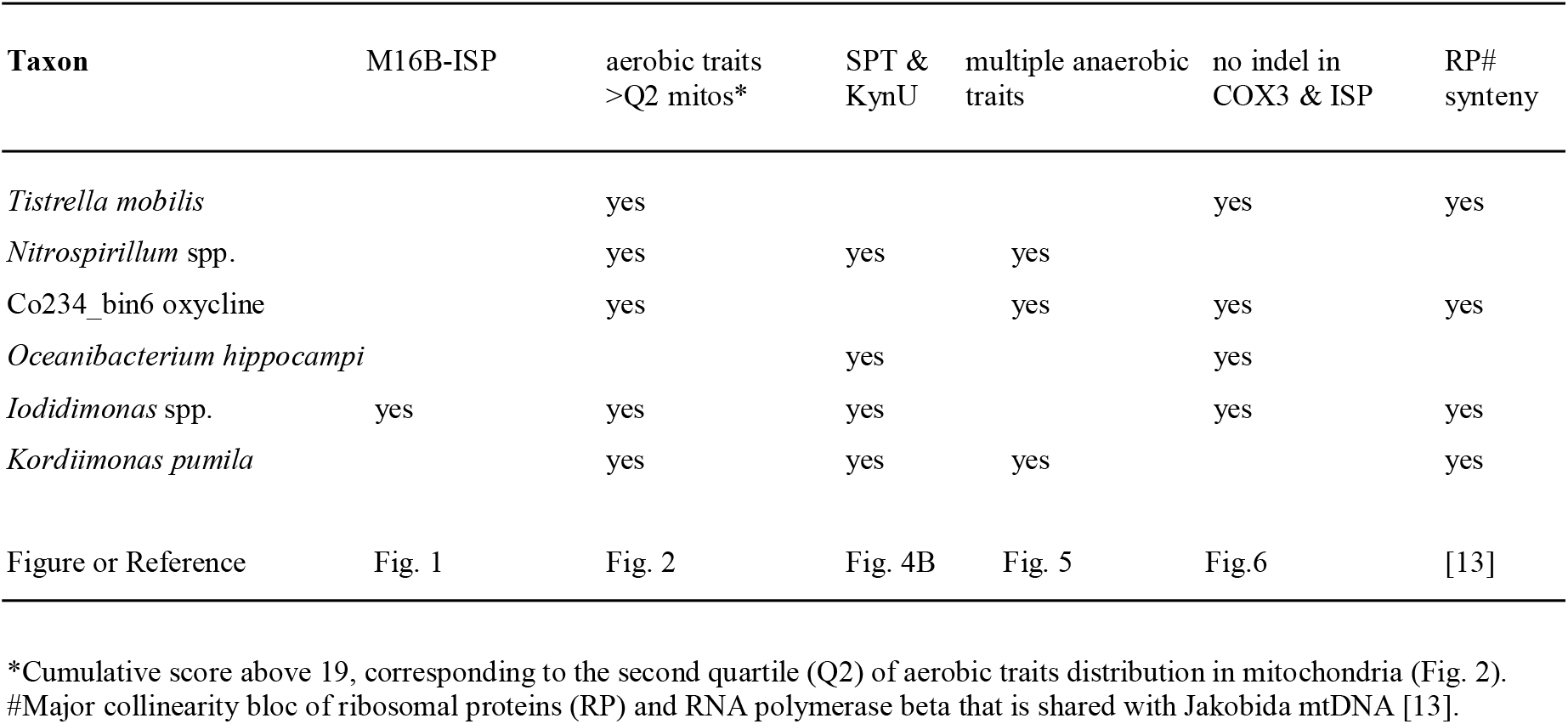
Selection criteria to evaluate the probability of alphaproteobacteria to be close to protomitochondria.

| Taxon | M16B-ISP | aerobic traits >Q2 mitos* | SPT & KynU | multiple anaerobic traits | no indel in COX3 & ISP | RP# synteny |
| --- | --- | --- | --- | --- | --- | --- |
| *Tistrella mobilis* |  | yes |  |  | yes | yes |
| *Nitrospirillum* spp. |  | yes | yes | yes |  |  |
| Co234_bin6 oxycline |  | yes |  | yes | yes | yes |
| *Oceanibacterium hippocampi* |  |  | yes |  | yes |  |
| *Iodidimonas* spp. | yes | yes | yes |  | yes | yes |
| *Kordiimonas pumila* |  | yes | yes | yes |  | yes |
| Figure or Reference | Fig. 1 | Fig. 2 | Fig. 4B | Fig. 5 | Fig.6 | [13] |

\*Cumulative score above 19, corresponding to the second quartile (Q2) of aerobic traits distribution in mitochondria (Fig. 2).
#Major collinearity bloc of ribosomal proteins (RP) and RNA polymerase beta that is shared with Jakobida mtDNA [13].

## CONCLUSION

Our triangulation strategy of sequential selection of extant bacteria with multiple approaches of different nature and bias (Tables 1 and 2) converges in indicating that the closest living bacterial relatives of protomitochondria may be present in Iodidimonadales and close lineages lying in the middle of alphaproteobacteria phylogeny. These bacteria are related to poorly characterized, earlier branching MAGs living in niches comparable to those present in proteorozoic oceans [10,35,46]. In turn, they are phylogenetically close to various MAGs of New clades thriving in marine zones with gradient oxygen [2,46] - the kind of environment that may be nearest to that pervading Proterozoic oceans when protomitochondria evolved [10,35]. Therefore, our novel insights dovetail the emerging picture that a facultative aerobe was the likely bacterial ancestor of protomitochondria [10], pinpointing to *Iodidimonas* spp. as its extant relatives.

## Supporting information

This file contains Supplementary Figures S1-S4 and Supplementary Table S1

## Acknowledgements

We thank Diego Gonzalez-Halphen, Esperanza Martinez-Romero, Miguel Angel Cevallos, Maria Maldonado, Seigo Amachi, Carolina Martinez-Gutierrez, Lars Hederstedt and Bill Martin for discussion and feed-back. We also thank Luis Lozano for technical assistance. O.G. acknowledges the support by grants from Dirección General de Asuntos del Personal Académico/Universida Nacional Autónoma de México (IN201120) and Consejo Nacional de Ciencia y Tecnología de México (118 in Investigación en Fronteras de la Ciencia).

## Data Availability

Supplementary Table S2 to S5 have ben deposited in figshare, 10.6084/m9.figshare.19768546. The repository of re-annotated genomes, the script in R to make searches in these genomes and alignments of respiratory proteins for analyzing INDEls will be made available upon request to the corresponding Author.

## Materials and Methods

The aim of the project leading to this paper was to unveil the metabolic ancestry of protomitochondria. To achieve this, we implemented a triangulation strategy by integrating evidence across phylogenomic and molecular analytical approaches that have different sources of bias [12,45] (Table 1). We applied these approaches to currently available alphaproteobacteria, lithotrophic gammaproteobacteria and eukaryotes with aerobic mitochondria. We excluded other bacteria because they do not have the subtype of COX operon that is characteristic of alphaproteobacteria and, in part, of mitochondria from early branching eukaryotes (Fig. 1, cf. [11]). Recognizing that it is indispensable to consider the whole spectrum of alphaproteobacteria diversity to evaluate their possible relationships with protomitochondria, we built an in-house repository of about 400 genomes representing all the major lineages of the class [2] plus the group of MarineProteo1 [1]. The genomes were systematically re-annotated for all the genes encoding proteins involved in oxidative phosphorylation and other metabolic pathways examined. Iterative PSI-BLAST searches of representative proteins [2,43] were carried out to evaluate their completeness and re-annotation congruity.

### Analysis of the aerobic traits

We have focused on the cytochrome part of the respiratory chain to specifically evaluate the aerobic traits shared by mitochondria and bacteria: cytochrome *c* and its reductase, the *bc*_1_ complex or complex III, and its oxidase, COX or complex IV. We additionally considered the alternative oxidase, AOX [47], which is the only other terminal oxidase in eukaryotes. A script in R [2] was built to systematically conduct and automate searches of the genes of these respiratory complexes, as well as those for the biogenesis of cytochrome *c*, heme A and copper cofactors. We thus constructed a preliminary presence/absence table for these proteins, assigning a value of 1 for complete and 0.5 for incomplete sequences. A −1 penalty was given for the absence of COX1 and COX2 of A1 type COX (Box1), the type of catalytic subunits of mitochondrial complex IV which must be present in possible ancestors of mitochondria [11]. The other proteins encoded by the genes of COX operon A1 type subtype b [11] were considered as traits except CtaB (Cox10), which is not encoded in any mtDNA nor is part of plant or protist complex IV. The catalytic subunits of complex III were also considered traits, together with the Cbp3 chaperone for the *bc*_1_ complex [23] (Box1). At difference with previous presence-absence analyses [2,7,11,13,14], we assigned a premium score of 2 for the following syntenic associations: i) COX1-COX2, forming a collinearity in the mtDNA of *Andalucia* [13] and marine members of the TSAR super-group [48] – they are also fused together in the mtDNA of Amoebozoa [11]; ii) Cox11 and COX3, which are syntenic in the mtDNA of *Andalucia* [13] and a few other early branching Eukaryotes (Supplementary Fig. S1A); M16B and ISP, which form a rare synteny (Fig. 1), possibly reflecting an ancestral genomic association before the migration of both genes to the nucleus (see text). We assigned a value of 1 to the genes for M16B.016 peptidases closest to eukaryotic MPP [22], identified by specific signatures in the aligned sequences, whenever such genes were associated to the COX operon (Fig.1). Additionally, we assigned a premium of 1 whenever the gene for CcmF, the critical heme lyase of System I for cytochrome *c* biogenesis in bacteria (Ccm [49]), was surrounded by at least two other Ccm genes. Multiple genes for the same trait within a genome were generally quantified with a numeral equivalent to such genes. Bacterial SCO and eukaryotic AOX showed such relatively uncommon situations (Supplementary Table S2). Conversely, multiple genes for M16A and M16C peptidases frequently occur in eukaryotic genomes ([26] and https://www.ebi.ac.uk/merops/cgi-bin/famsum?family=M16 - accessed on 11 April 2022). In such cases, we capped the maximal numeral at 5 (Box1). The last two traits listed in Box1 correspond to signatures that are present in two catalytic proteins that are indispensable for aerobic metabolism: COX1 and ISP. These were chosen among various signatures because they are common in taxa with reduced genomes such as those of Rickettsiales, which are more prone to differential gene loss than other lineages [7]. COX1 proteins of Archaeplastida and several alphaproteobacterial lineages contain a cysteine (Cys) lying in a conserved region at the negative side of the membrane (cytoplasm in bacteria or matrix in mitochondria), between the last transmembrane segment and the C terminus. This region is critical for the assembly of Complex IV in yeast and presumably other eukaryotes [25]. The signature in ISP was considered as the absence of its major INDEls, one just before the conserved ligand blocs for the iron-sulfur cluster and the other in the middle of the same ligand blocs [11], which occurs in mitochondria and a limited number of alphaproteobacteria including Rickettsiales (Supplementary Table S4). In sum, the cumulative score of the 18 aerobic traits we have analysed here provides a multi-layered evaluation of bacterial and eukaryotic metabolism never considered before.

### Taxonomic sampling of mitochondria and grouping of alphaproteobacteria in 15 lineages

We have dedicated extensive efforts to thoroughly deal with the issue of taxonomic sampling, most critical for evaluating the bacterial origin of mitochondria [1–8,30]. The initial efforts focused on the selection of eukaryotic taxa that have aerobic mitochondria and can be considered early branching [50]. The selection of the aerobic traits (Box 1) guided the choice of thes eukaryotic taxa, which were selected on the basis of the presence of at least one Ccm gene for cytochrome *c* biogenesis, because the Ccm system has been vertically inherited from alphaproteobacteria [13,14,49]. CcmF and other components of the Ccm system are encoded in the mtDNA of a variety of eukaryotic taxa including plants, Jakobida [13,15], *Diphylleia* [14], *Palpitomons* [51], Cryptista-related such as *Hemiarma* [52], Malawimonadida, Cyanidiales of Rhodophyta [53] and also Ciliophora such as *Tetrahymena* [54]. Ciliophora are at odds with the rest of the TSAR super-group to which they belong [50], which generally has System III for cytochrome *c* biogenesis [49,52]. System III consists of a single heme lyase protein without bacterial precedents, therefore constituting a eukaryotic innovation [49,52]. For this reason, we have excluded the eukaryotic groups that have System III and its possible ancestor System V of Euglenozoa [55], except for lineages that are related to those containing the Ccm system (Cryptista, Haptista, Chlorophyta and Rhodophyta [52]). We additionally excluded Ciliophora for the highly derived nature of their genomes [16,54] and Malawimonadida because they lack nuclear genes. After detailed analysis, in part used for other aspects of this work, we also excluded Rhodelphida [53] and Picozoa [56], because their genome is incomplete and does not encode the Ccm system. Considering the limited number of nuclear genomes that are currently available for Discoba and Archaeplastida other than plants, we settled to analyze a set of about 30 Eukaryotes to represent aerobic mitochondria (Supplementary Table S2). Permutations of some taxa of this set with others including yeast, *Thecamonas* and Amoebozoa such as *Dyctiostelium* - which we used in some phylogenetic analyses - did not significantly alter the distribution of the aerobic traits in mitochondria and their cumulative score, which maintained a median value of 21 (Fig. 2). Next, we endeavoured to produce a thorough selection of alphaproteobacteria representing all the major lineages of the class with their specific metabolic traits. Our previous taxonomic analysis [2] guided the selection of 15 separate lineages of alphaproteobacteria, spanning from early-branching Rickettsiales to late-branching Rhodobacterales. Intermediate lineages also corresponded to orders in the current taxonomy of alphaproteobacteria (https://www.ncbi.nlm.nih.gov/Taxonomy/Browser/www.tax.cgi?id=28211, accessed on 11 April 2022), which essentially derives from a recent re-classification [57]. However, we considered as separate lineages two groups that most likely belong to the Rhodospirillales super-order [2] due to phylogenomic and other observations, as follows. HIMB59 and related marine MAGs were clustered together in the lineage called ‘MarineAlpha’, which includes other marine bacteria with similarly AT-rich genomes, in particular those classified under the TMED109 and TMED127 orders in GTDB taxonomy [58]. Holosporales, which generally cluster with clades of Rhodospirillales after correcting for their AT-rich genomes [7,8], were considered an intermediate lineage between Rickettsiales and Rhodospirillales, a placement sustained by their low cumulative scores of aerobic traits (Fig. 2). We used around 60 taxa to encompass the wide genomic diversity of the Rhodospirillales super-order, including at least two taxa for each major subdivision [2]. We routinely kept the recently identified New clades of marine MAGs as a separate lineage from related Sneathiellales (including also Minwuiales) because of the diversity in their bioenergetic traits [2]. We also kept the lineage of Iodidimonadales separate from other SERIK groups [2] essentially because of their unique synteny of M16B-ISP (Fig. 1 and Supplementary Fig. S1). To increase the genomic diversity of Iodidimonadales, we included two alphaproteobacterial MAGs found in an anoxic hydrothermal niche [46], proteins of which cluster together with those of *Iodidimonas* in phylogenetic trees (for example COX3 in Supplementary Fig. S4B). Conversely, we merged the order of Rhodothalassiales and Kordiimonadales, as well that of Zavarziniales with Emcibacterales, by considering common features such as number of INDELs in respiratory proteins. Representatives of all major families were included to encompass the genomic diversity of Rhizobiales [2,57]. In contrast, only the deepest branching and major genera were included to represent the limited taxonomic diversity of the lineages of Pelagibacterales, Sphingomonadales, Caulobacterales and Rhodobacterales. The Pelagibacterales lineage was placed just before the Rhizobiales lineage in accordance with recent phylogenetic analyses [2,8]. We did not consider the orders of Parvularculales and Maricaulales, because they contain late branching taxa that often cluster with representatives of the established orders of Caulobacterales or Rhodobacterales [2,57]. In all cases, the composition of the alphaproteobacterial lineages was balanced to provide a good match in relative frequency of bioenergetic traits with respect to the complete set of available genomes, as previously verified for NosZ [2]. Although we favored the inclusion of cultivated taxa with complete genome, in several cases this was not possible because the lineage or its taxonomic divisions include only MAGs, as in the case of MarineProteo1 [1,8]. In such lineages, we considered MAGs with most complete genomes, as deduced from the GTDB database or our own evaluations conducted as described earlier [2]. MarineProteo1_Bin1 was kept despite its poor coverage because of its prototypic position in the MarineProteo1 lineage [1,8]. In sum, the lineage-dependent plots of various traits we present in this work reflect divergence time along the x axis, following the consistent branching order of alphaproteobacteria in phylogenetic trees [2].

### Distribution of the traits for ceramide and kynurenine biosynthesis

The second approach we used in (Table 1) followed a conventional absence-presence analysis (Supplementary Table S3). We first conducted PSI-BLAST searches of putative homologs of *Caulobacter* SPT [27,29] in all the genomes of our in-house repository, plus those of closely related taxa. An Evalue cut-off of 5e^-68^ was generally sufficient to differentiate genuine SPT homologs from related α-oxo-amine synthases such as 5-aminolevulinate synthase. The genomic regions containing the gene encoding the identified SPT homologs were then carefully inspected to verify the presence of the four-genes operon that is common in ceramide-synthesizing bacteria [29] (Fig. 3A), and the eventual vicinity of other genes required for sphingolipid biosynthesis [27]. Each protein of the operon was then analyzed in detailed alignments to identify its biochemical nature (Supplementary Table S3). Various bacterial SPT proteins were then used in PSI-BLAST searches extended to eukaryotes, focusing on the species selected for the comparative analysis of the aerobic traits (Supplementary Table S2). We considered both the active and inactive isoform of eukaryotic SPT, which likely derive from a process of duplication and differentiation of the single *spt* gene of bacteria [28] (Fig. 3B). An equivalent strategy was used for identifying bacterial and eukaryotic homologs of *Pseudomonas* kynureninase, encoded by the *kynU* gene which is often adjacent to *kynA* and other genes for components of the kynurenine pathway [33,34].

### Analysis of anaerobic traits

We have expanded the analysis of the kynurenine pathway to evaluate whether such pathway might be involved in *de novo* biosynthesis of RQ [34] in alphaproteobacteria. We cross-checked this analysis with the evaluation of the taxonomic distribution of the *rquA* gene discovered in *R. rubrum*, which enables the conversion of RQ from pre-formed Q in this and other bacteria [36]. We found the distribution of the *rquA* gene to be mutually exclusive with the combination of two genes for UbiA with the *kynU* gene adjacent to other genes for the kynurenine pathway (see text, cf. Fig. 5). This mutual exclusivity has been verified in the genomes of the genus *Rhodoferax* known to contain RQ [60]. For example, *Rhodoferax fermentans* has the *rquA* gene (protein accession WP_078364963) but not the *kynU* gene, whereas *Rhodoferax sediminis* has the *kynU* gene for a functional kynureninase (protein accession WP_142820272) but no homolog of the *rquA* gene.

The presence of the *ubiTUV* triad of genes required for the anaerobic biosynthesis of Q [37] was identified by the genomic collinearity of these genes and protein alignments defining the distinctive signatures of the coded proteins [2], including the conserved Cys ligands of the 4Fe4S cluster [37,38]. Blast searches extended to eukaryotes identified diverse paralogs with the same U32 peptidase domain as UbiU [37], including composite proteins showing the fusion of this domain with the DUF3656 domain as in the protein coded by *E.coli rlhA* [38,61], often in combination with as similar, UbiV-like domain fused at the C-terminus. Most frequently, such proteins are found among members of the Sar super-group [50] and Chlorophyta; however, similar proteins are present in Magnetococcales and other deep branching alphaproteobacteria. Eukaryotic protein hits found among invertebrates were identified instead as likely bacterial contaminants on the basis of dedicated blast searches.

Identification of the different types of OFORs was undertaken following the molecular analysis described in Ref. [39]. Their presence was quantified assigning a value of 1 to each monomeric or heterodimeric type, and of 0.5 whenever a coded protein was incomplete. OFORs involved in the biosynthesis of photosynthetic pigments were ignored.

### Phylogenetic and molecular analysis of proteins

Phylogenetic analysis was conducted with large alignments of diverse protein sequences [2,43]. Such alignments were manually implemented after rounds of automated alignment with the MUSCLE program within the MEGA software (versions 5 and X [2,43]). Phylogenetic analysis was undertaken with Maximum Likelihood (ML) inference using the program IQ-Tree and Bayesian inference using the BEAST program, as previously described [2,43]. The LG models was most frequently used. Analysis of the INDELs present in the catalytic subunits of complex III and IV was conducted in enlarged versions of the same alignments used for phylogenetic analysis. Such alignments included proteins from all the bacterial and eukaryotic taxa that we examined for the aerobic traits. Short deletions or insertions that were species-specific were ignored, while INDELs at least three amino acids long and shared by at least three bacterial lineages were considered for the analysis (Supplementary Table S4). Alignments of the NuoL/ND5 and NuoD/Nad7 proteins were built and analyzed in the same way. A total of 21 conserved INDELs, predominantly constituted by insertions not shared with mitochondrial homologs (Supplementary Fig. S3A), were identified in respiratory proteins using the systematic application of the above criterion. Their distribution was evaluated in the same lineage-specific way as for the aerobic traits. Of note, several other proteins were not considered because they presented variable INDELs in eukaryotic sequences vs. their bacterial homologs, as for example in CtaG_Cox11.

### Statistical analysis

Statistical analysis was conducted with the 99% Confidence Interval of the T test [59]. In particular, we conducted independent t-tests to verify the two-tailed hypothesis that the cumulative scores of the 18 aerobic traits (Box 1) for each lineage significantly differed from mitochondria. To guard against inflated type 1 errors (i.e., false positives) from multiple testing, these analyses were conducted at the more stringent 1% significance level (alpha=0.01). All analyses were run in R (version 4.1.0) using the stats package.

## Supplementary Material

The Supplementary Material of this paper includes four Supplementary Figures and five Supplementary Tables.

**Supplementary Figure S1.** A. Synteny COX11-COX3 in the mtDNA of early-branching Eukaryotes, with progressive distance by additional genes. B. Phylogenetic tree of M16B and MPP proteins. The ML tree was reconstructed from an alignment of equal number (33) of bacterial M16B and eukaryotic MPP proteins, using the best fit model LG obtained with the IQ-Tree web server system [43]. Note that the isolated M16B of *Rickettsia conorii* has been experimentally found to cleave mitochondrial pre-sequences as eukaryotic MPP proteins [21]. The outgroup proteins belong to the UPB group that forms part of a syntenic diad of M16B metallopeptidases, which are distinct from the M16.019 group closest to eukaryotic MPPbeta [22]. Similar trees were obtained with enlarged alignments and the EX_EHO mixture model.

**Supplementary Figure S2.** M16B-ISP collinearity in the genome of some alphaproteobacteria.

**Supplementary Figure S3.** A. The lineage-specific distribution of the sums of all 2-oxoacid ferredoxin oxido reductases (OFORs) - not related to photosynthesis - is represented in mean plus SD values (cf. Figs. 5 and 6). B. Comparison of the distribution of cumulative scores for aerobic traits (cf. Fig. 2) and the sums of OFORs anaerobic traits along the lineages of alphaproteobacteria (Supplementary Table S2, rightmost column). *The median values were subtracted by 6, the median of aerobic scores for gammaproteobacteria (Fig. 2).

**Supplementary Figure S4.** A. INDELs in COX3 sequences. The number before and after the INDELs correspond to the position in the alignment of diverse bacterial (top) and mitochondrial (bottom) COX3 proteins. B. Mapping of the INDELs of COX3 along a ML tree of the protein. The ML tree was reconstructed from an alignment of 140 COX3 sequences equivalent to that used in part A. It is representative of 20 similar trees obtained with different models and slightly different combinations of aligned proteins.

**Supplementary Table S1.** MPP and related proteins in various eukaryotic taxa. Support for Fig. 1, shown directly in the Supplementary Material .pdf file.

Other Supplementary Tables have been inserted in a datasheet deposited in figshare with the reference: 10.6084/m9.figshare.19768546

This datasheet contains the Supplementary Tables listed below, which are separate sheets of a single .excel file deposited in figshare to complement the data and analyses of the manuscript entitled: On the bacterial ancestry of mitochondria: new insights with triangulated approaches

**Supplementary Table S2.** Tabulation of 18 aerobic traits and their cumulative score in 15 alphaproteobacterial lineages plus a set of aerobic mitochondria from eukaryotes. The rightmost column shows the sum values for the 2-oxoacid ferredoxin oxidoreductases (OFORs) considered as traits of anaerobic metabolism.

**Supplementary Table S2B.** Statistical results of the T test on the lineage-specific scores of aerobic traits listed in Table S2.

**Supplementary Table S3.** Distribution of *spt* (the gene for serine-palmitoyl transferases) and allied genes for ceramide biosynthesis, as well as of *kynU* (gene for kynurenonase) and allied genes of the kynurenine pathway among 15 lineages of alphaproteobacteria. When the *kynA* gene for tryptophan di-oxygenase gene lies in the same genomic region of *kynU* it is annotated as 1.

**Supplementary Table S4**. Distribution of conserved INDELs in the indicated respiratory proteins.

## References

1. Martijn J, Vosseberg J, Guy L, Offre P, Ettema TJG. 2018. Deep mitochondrial origin outside the sampled alphaproteobacteria. Nature. 55:101–105. doi: 10.1038/s41586-018-0059-5.

2. Cevallos MA, Degli Esposti M. 2022. New Alphaproteobacteria Thrive in the Depths of the Ocean with Oxygen Gradient. Microorganisms. 10:455. doi: 10.3390/microorganisms10020455

3. Gray MW, Burger G, Lang BF. 1999. Mitochondrial evolution. Science. 283:1476–1481. doi: 10.1126/science.283.5407.1476

4. Ferla MP, Thrash JC, Giovannoni SJ, Patrick WM. 2013. New rRNA gene-based phylogenies of the Alphaproteobacteria provide perspective on major groups, mitochondrial ancestry and phylogenetic instability. PLoS One. 8:e83383. doi: 10.1371/journal.pone.0083383. PMID: 24349502; PMCID: PMC3859672

5. Esser C, Ahmadinejad N, Wiegand C, Rotte C, Sebastiani F, Gelius-Dietrich G, Henze K, Kretschmann E, Richly E, Leister D, Bryant D, Steel MA, Lockhart PJ, Penny D, Martin W. 2004. A genome phylogeny for mitochondria among alphaproteobacteria and a predominantly eubacterial ancestry of yeast nuclear genes. Mol Biol Evol. 21:1643–1660. doi: 10.1093/molbev/msh160

6. Nagies FSP, Brueckner J, Tria FDK, Martin WF. 2020. A spectrum of verticality across genes. PLoS Genet. 16:e1009200. doi: 10.1371/journal.pgen.1009200.

7. Fan L, Wu D, Goremykin V, Xiao J, Xu Y, Garg S, Zhang C, Martin WF, Zhu R. 2020. Phylogenetic analyses with systematic taxon sampling show that mitochondria branch within Alphaproteobacteria. Nat Ecol Evol. 4:1213–1219. doi: 10.1038/s41559-020-1239-x.

8. Muñoz-Gómez SA, Susko E, Williamson K, Eme L, Slamovits CH, Moreira D, López-García P, Roger AJ. Site-and-branch-heterogeneous analyses of an expanded dataset favour mitochondria as sister to known Alphaproteobacteria. Nat Ecol Evol. 2022 Mar;6(3):253–262. doi: 10.1038/s41559-021-01638-2

9. Imachi H, Nobu MK, Nakahara N, Morono Y, Ogawara M, Takaki Y, Takano Y, Uematsu K, Ikuta T, Ito M, Matsui Y, Miyazaki M, Murata K, Saito Y, Sakai S, Song C, Tasumi E, Yamanaka Y, Yamaguchi T, Kamagata Y, Tamaki H, Takai K. 2020. Isolation of an archaeon at the prokaryote-eukaryote interface. Nature. 577:519–525. doi: 10.1038/s41586-019-1916

10. Mills DB, Boyle RA, Daines SJ, Sperling EA, Pisani D, Donoghue PCJ, Lenton TM. 2022. Eukaryogenesis and oxygen in Earth history. Nat Ecol Evol. 6:520–532. doi: 10.1038/s41559-022-01733-y.

11. Degli Esposti M, Chouaia B, Comandatore F, Crotti E, Sassera D, Lievens PM, Daffonchio D, Bandi C. 2014. Evolution of mitochondria reconstructed from the energy metabolism of living bacteria. PLoS One. 9:e96566. doi: 10.1371/journal.pone.0096566

12. Munafò MR, Davey Smith G. 2018. Robust research needs many lines of evidence. Nature. 553:399–401. doi: 10.1038/d41586-018-01023-3

13. Burger G, Gray MW, Forget L, Lang BF. 2013. Strikingly bacteria-like and gene-rich mitochondrial genomes throughout jakobid protists. Genome Biol. Evol. 5:418–438. doi: 10.1093/gbe/evt008.

14. Kamikawa R, Shiratori T, Ishida K, Miyashita H, Roger AJ. Group II Intron-Mediated Trans-Splicing in the Gene-Rich Mitochondrial Genome of an Enigmatic Eukaryote, Diphylleia rotans. Genome Biol Evol. 2016 Feb 1;8(2):458–66. doi: 10.1093/gbe/evw011

15. Yabuki A, Gyaltshen Y, Heiss AA, Fujikura K, Kim E. 2018. Ophirina amphinema n. gen., n. sp., a new deeply branching discobid with phylogenetic affinity to jakobids. Sci Reports. 8:1–14. doi: 10.1038/s41598-018-34504-6.

16. Zhou L, Maldonado M, Padavannil A, Guo F, Letts JA. 2022. Structures of Tetrahymena’s respiratory chain reveal the diversity of eukaryotic core metabolism. Science. 10.1126/science.abn7747.

17. Zhang Z, Huang L, Shulmeister VM, Chi YI, Kim KK, Hung LW, Crofts AR, Berry EA, Kim SH. 1998. Electron transfer by domain movement in cytochrome bc1. Nature. 392:677–684. doi: 10.1038/33612

18. Braun HP, Schmitz UK. 1995. Are the ‘core’ proteins of the mitochondrial bc1 complex evolutionary relics of a processing protease? Trends Biochem Sci. 20:171–175. doi: 10.1016/s0968-0004(00)88999-9

19. Maldonado M, Guo F, Letts JA. 2021. Atomic structures of respiratory complex III2, complex IV, and supercomplex III2-IV from vascular plants. Elife. 10:e62047. doi: 10.7554/eLife.62047

20. Iwata S, Lee JW, Okada K, Lee JK, Iwata M, Rasmussen B, Link TA, Ramaswamy S, Jap BK. 1998. Complete structure of the 11-subunit bovine mitochondrial cytochrome bc1 complex. Science. 281:64–71. doi: 10.1126/science.281.5373.64

21. Kitada S, Uchiyama T, Funatsu T, Kitada Y, Ogishima T, Ito A. 2007. A protein from a parasitic microorganism, Rickettsia prowazekii, can cleave the signal sequences of proteins targeting mitochondria. J Bacteriol. 189:844–850

22. Aleshin AE, Gramatikova S, Hura GL, Bobkov A, Strongin AY, Stec B, Tainer JA, Liddington RC, Smith JW. 2009. Crystal and solution structures of a prokaryotic M16B peptidase: an open and shut case. Structure. 17:1465–1475. doi: 10.1016/j.str.2009.09.009

23. Barros MH, McStay GP. 2020. Modular biogenesis of mitochondrial respiratory complexes. Mitochondrion. 50:94–114. doi: 10.1016/j.mito.2019.10.008

24. Kubo N, Arimura S, Tsutsumi N, Kadowaki K, Hirai M. 2006. Isolation and characterization of the pea cytochrome c oxidase Vb gene. Genome. 49:1481–1489. doi: 10.1139/g06-105

25. García-Villegas R, Camacho-Villasana Y, Shingú-Vázquez MÁ, Cabrera-Orefice A, Uribe-Carvajal S, Fox TD, Pérez-Martínez X. 2017. The Cox1 C-terminal domain is a central regulator of cytochrome c oxidase biogenesis in yeast mitochondria. J Biol Chem. 292:10912–10925. doi: 10.1074/jbc.M116.773077

26. Grinter R, Hay ID, Song J, Wang J, Teng D, Dhanesakaran V, Wilksch JJ, Davies MR, Littler D, Beckham SA, Henderson IR, Strugnell RA, Dougan G, Lithgow T. 2018. FusC, a member of the M16 protease family acquired by bacteria for iron piracy against plants. PLoS Biol. 16:e2006026. doi: 10.1371/journal.pbio.2006026

27. Olea-Ozuna RJ, Poggio S, EdBergström, Quiroz-Rocha E, García-Soriano DA, Sahonero-Canavesi DX, Padilla-Gómez J, Martínez-Aguilar L, López-Lara IM, Thomas-Oates J, Geiger O. 2021. Five structural genes required for ceramide synthesis in Caulobacter and for bacterial survival. Environ Microbiol. 23:143–159. doi: 10.1111/1462-2920.15280

28. Ikushiro H, Islam MM, Tojo H, Hayashi H. 2007. Molecular characterization of membrane-associated soluble serine palmitoyltransferases from Sphingobacterium multivorum and Bdellovibrio stolpii. J Bacteriol. 189:5749–5761. doi: 10.1128/JB.00194-07

29. Padilla-Gómez J, Olea-Ozuna RJ, Contreras-Martínez S, Morales-Tarré O, García-Soriano DA, Sahonero-Canavesi DX, Poggio S, Encarnación S, López-Lara IM, Geiger O. 2022. Specialized acyl carrier protein used by serine palmitoyltransferase to synthesize sphingolipids in Rhodobacteria. J Biol Chem., in press.

30. Degli Esposti M. 2014. Bioenergetic evolution in proteobacteria and mitochondria. Genome Biol Evol. 6:3238–51. doi: 10.1093/gbe/evu257

31. Muñoz-Gómez SA, Wideman JG, Roger AJ, Slamovits CH. 2017. The Origin of Mitochondrial Cristae from Alphaproteobacteria. Mol Biol Evol. 34:943–956. doi: 10.1093/molbev/msw298

32. Alonso A, Goñi FM. The Physical Properties of Ceramides in Membranes. Annu Rev Biophys. 2018 May 20;47:633–654. doi: 10.1146/annurev-biophys-070317-033309. Epub 2018 Apr 4. PMID: 29618220.

33. Phillips RS. Structure, mechanism, and substrate specificity of kynureninase. 2011. Biochim Biophys Acta. 1814:1481–1488. doi: 10.1016/j.bbapap.2010.12.003

34. Salinas G, Langelaan DN, Shepherd JN. 2020. Rhodoquinone in bacteria and animals: Two distinct pathways for biosynthesis of this key electron transporter used in anaerobic bioenergetics. Biochim Biophys Acta Bioenerg. 1861:148278. doi: 10.1016/j.bbabio.2020.148278

35. Müller M, Mentel M, van Hellemond JJ, Henze K, Woehle C, Gould SB, Yu RY, van der Giezen M, Tielens AG, Martin WF. 2012. Biochemistry and evolution of anaerobic energy metabolism in eukaryotes. Microbiol Mol Biol Rev. 76:444–495. doi: 10.1128/MMBR.05024-11

36. Stairs CW, Eme L, Muñoz-Gómez SA, Cohen A, Dellaire G, Shepherd JN, Fawcett JP, Roger AJ. 2018. Microbial eukaryotes have adapted to hypoxia by horizontal acquisitions of a gene involved in rhodoquinone biosynthesis. Elife. 7:e34292. doi: 10.7554/eLife.34292

37. Pelosi L, Vo CD, Abby SS, Loiseau L, Rascalou B, Hajj Chehade M, Faivre B, Goussé M, Chenal C, Touati N, Binet L, Cornu D, Fyfe CD, Fontecave M, Barras F, Lombard M, Pierrel F. 2019. Ubiquinone Biosynthesis over the Entire O2 Range: Characterization of a Conserved O2-Independent Pathway. mBio. 10:e01319–19. doi: 10.1128/mBio.01319-19

38. Abby SS, Kazemzadeh K, Vragniau C, Pelosi L, Pierrel F. 2020. Advances in bacterial pathways for the biosynthesis of ubiquinone. Biochim Biophys Acta Bioenerg. 1861:148259. doi: 10.1016/j.bbabio.2020.148259

39. Chen PY, Li B, Drennan CL, Elliott SJ. 2019. A reverse TCA cycle 2-oxoacid:ferredoxin oxidoreductase that makes C-C bonds from CO2. Joule. 3:595–611. doi: 10.1016/j.joule.2018.12.006

40. Gupta RS. 2005. Protein signatures distinctive of alpha proteobacteria and its subgroups and a model for alphaproteobacterial evolution. Crit Rev Microbiol. 31:101–135. doi: 10.1080/10408410590922393

41. Lavrov DV. 2007. Key transitions in animal evolution: a mitochondrial DNA perspective. Integr Comp Biol. 47:734–743. doi: 10.1093/icb/icm045

42. Ajawatanawong P, Baldauf SL. 2013. Evolution of protein indels in plants, animals and fungi. BMC Evol Biol. 13:140. doi: 10.1186/1471-2148-13-140

43. Degli Esposti M, Moya-Beltrán A, Quatrini R, Hederstedt L. 2021. Respiratory Heme A-Containing Oxidases Originated in the Ancestors of Iron-Oxidizing Bacteria. Front Microbiol. 12:664216. doi: 10.3389/fmicb.2021.664216

44. Giovannoni SJ, Tripp HJ, Givan S, Podar M, Vergin KL, Baptista D, Bibbs L, Eads J, Richardson TH, Noordewier M, Rappé MS, Short JM, Carrington JC, Mathur EJ. 2005. Genome streamlining in a cosmopolitan oceanic bacterium. Science. 309:1242–1245. doi: 10.1126/science.1114057

45. Lawlor DA, Tilling K, Davey Smith G. 2016. Triangulation in aetiological epidemiology. Int J Epidemiol. 45:1866–1886. doi: 10.1093/ije/dyw314

46. Ward LM, Idei A, Nakagawa M, Ueno Y, Fischer WW, McGlynn SE. 2019. Geochemical and Metagenomic Characterization of Jinata Onsen, a Proterozoic-Analog Hot Spring, Reveals Novel Microbial Diversity including Iron-Tolerant Phototrophs and Thermophilic Lithotrophs. Microbes Environ. 34:278–292. doi: 10.1264/jsme2.ME19017

47. Atteia A, van Lis R, van Hellemond JJ, Tielens AG, Martin W, Henze K. 2004. Identification of prokaryotic homologues indicates an endosymbiotic origin for the alternative oxidases of mitochondria (AOX) and chloroplasts (PTOX). Gene. 330:143–148. doi: 10.1016/j.gene.2004.01.015

48. Wideman JG, Monier A, Rodríguez-Martínez R, Leonard G, Cook E, Poirier C, Maguire F, Milner DS, Irwin NAT, Moore K, Santoro AE, Keeling PJ, Worden AZ, Richards TA. 2020. Unexpected mitochondrial genome diversity revealed by targeted single-cell genomics of heterotrophic flagellated protists. Nat Microbiol. 5:154–165. doi: 10.1038/s41564-019-0605-4

49. Verissimo AF, Daldal F. 2014. Cytochrome c biogenesis System I: an intricate process catalyzed by a maturase supercomplex? Biochim Biophys Acta. 1837:989–998. doi: 10.1016/j.bbabio.2014.03.003

50. Burki F, Roger AJ, Brown MW, Simpson AGB. Trends Ecol Evol. The New Tree of Eukaryotes. 2020 35:43–55. doi: 10.1016/j.tree.2019.08.008

51. Yabuki A, Inagaki Y, Ishida K. 2010. Palpitomonas bilix gen. et sp. nov.: A novel deep-branching heterotroph possibly related to Archaeplastida or Hacrobia. Protist. 161:523–538. doi: 10.1016/j.protis.2010.03.001

52. Nishimura Y, Kume K, Sonehara K, Tanifuji G, Shiratori T, Ishida KI, Tetsuo H, Inagaki Y, Ohkuma M. 2020. Mitochondrial genomes of Hemiarma marina and Leucocryptos marina revised the evolution of cytochrome c maturation in Cryptista. Frontiers Ecol Evol. 8:140. https://doi.org/10.3389/fevo.2020.00140

53. Gawryluk RMR, Tikhonenkov DV, Hehenberger E, Husnik F, Mylnikov AP, Keeling PJ. 2019. Non-photosynthetic predators are sister to red algae. Nature. 572:240–243. doi: 10.1038/s41586-019-1398-6

54. Brunk CF, Lee LC, Tran AB, Li J. 2003. Complete sequence of the mitochondrial genome of Tetrahymena thermophila and comparative methods for identifying highly divergent genes. Nucleic Acids Res. 31:1673–1682. doi: 10.1093/nar/gkg270

55. Belbelazi A, Neish R, Carr M, Mottram JC, Ginger ML. 2021 Divergent Cytochrome c Maturation System in Kinetoplastid Protists. mBio.12:e00166–21. doi: 10.1128/mBio.00166-21.

56. Schön ME, Zlatogursky VV, Singh RP, Poirier C, Wilken S, Mathur V, Strassert JFH, Pinhassi J, Worden AZ, Keeling PJ, Ettema TJG, Wideman JG, Burki F. 2021. Single cell genomics reveals plastid-lacking Picozoa are close relatives of red algae. Nat Commun. 12:6651. doi: 10.1038/s41467-021-26918-0

57. Hördt A, López MG, Meier-Kolthoff JP, Schleuning M, Weinhold LM, Tindall BJ, Gronow S, Kyrpides NC, Woyke T, Göker M. 2020. Analysis of 1,000+ Type-Strain Genomes Substantially Improves Taxonomic Classification of Alphaproteobacteria. Front Microbiol. 11:468. doi: 10.3389/fmicb.2020.00468

58. 2019. Chaumeil PA, Mussig AJ, Hugenholtz P, Parks DH. GTDB-Tk: a toolkit to classify genomes with the Genome Taxonomy Database. Bioinformatics. 36:1925–1927. doi: 10.1093/bioinformatics/btz848

59. Gerald B. 2018. A brief review of independent, dependent and one sample t-test. Int J Applied Mathe Theor Phys. 4:50–54.

60. Miyadera H, Hiraishi A, Miyoshi H, Sakamoto K, Mineki R, Murayama K, Nagashima KV, Matsuura K, Kojima S, Kita K. 2003. Complex II from phototrophic purple bacterium Rhodoferax fermentans displays rhodoquinol-fumarate reductase activity. Eur J Biochem. 270:1863–1874. doi: 10.1046/j.1432-1033.2003.03553.x

61. Kimura S, Sakai Y, Ishiguro K, Suzuki T. 2017. Biogenesis and iron-dependency of ribosomal RNA hydroxylation. Nucleic Acids Res. 45:12974–12986. doi: 10.1093/nar/gkx969 Additional References in Figure Legends

62. Gray MW. 2015. Mosaic nature of the mitochondrial proteome: Implications for the origin and evolution of mitochondria. Proc Natl Acad Sci U S A. 112:10133–10138. doi: 10.1073/pnas.1421379112

63. Iwata S. 1998. Structure and function of bacterial cytochrome c oxidase. J Biochem. 123:369–375. doi: 10.1093/oxfordjournals.jbchem.a021946

64. Iino T, Ohkuma M, Kamagata Y, Amachi S. 2016. Iodidimonas muriae gen. nov., sp. nov., an aerobic iodide-oxidizing bacterium isolated from brine of a natural gas and iodine recovery facility, and proposals of Iodidimonadaceae fam. nov., Iodidimonadales ord. nov., Emcibacteraceae fam. nov. and Emcibacterales ord. nov. Int J Syst Evol Microbiol. 66:5016–5022. doi: 10.1099/ijsem.0.001462

