## Supplementary material for "On the bacterial ancestry of mitochondria: new insights with triangulated approaches": This file contains Supplementary Figures S1-S4 and Supplementary Table S1

The Supplementary Material of this paper includes four Supplementary Figures and five Supplementary Tables.

Supplementary Figures S1 to S4 are presented below, followed by Supplementary Table S1.

The other Supplementary Tables form separate sheets of a separate .excel file.

**A.**

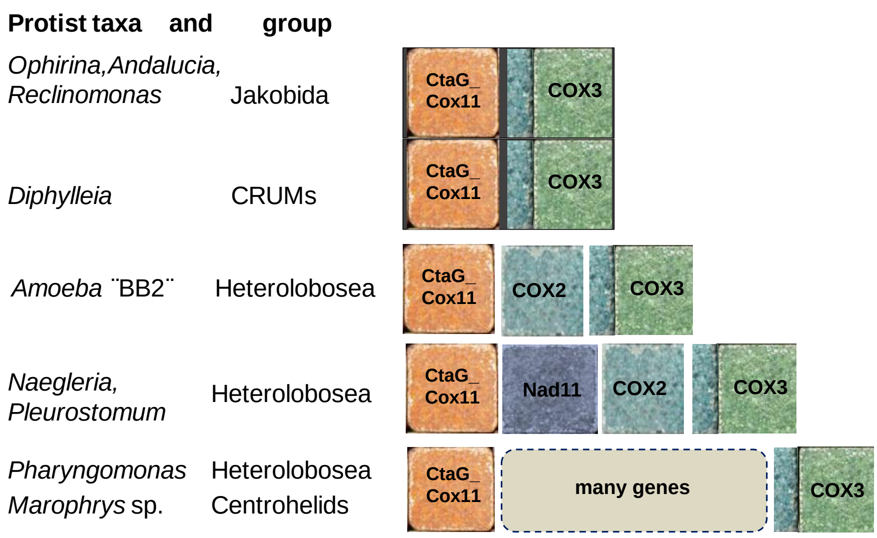

**B.**

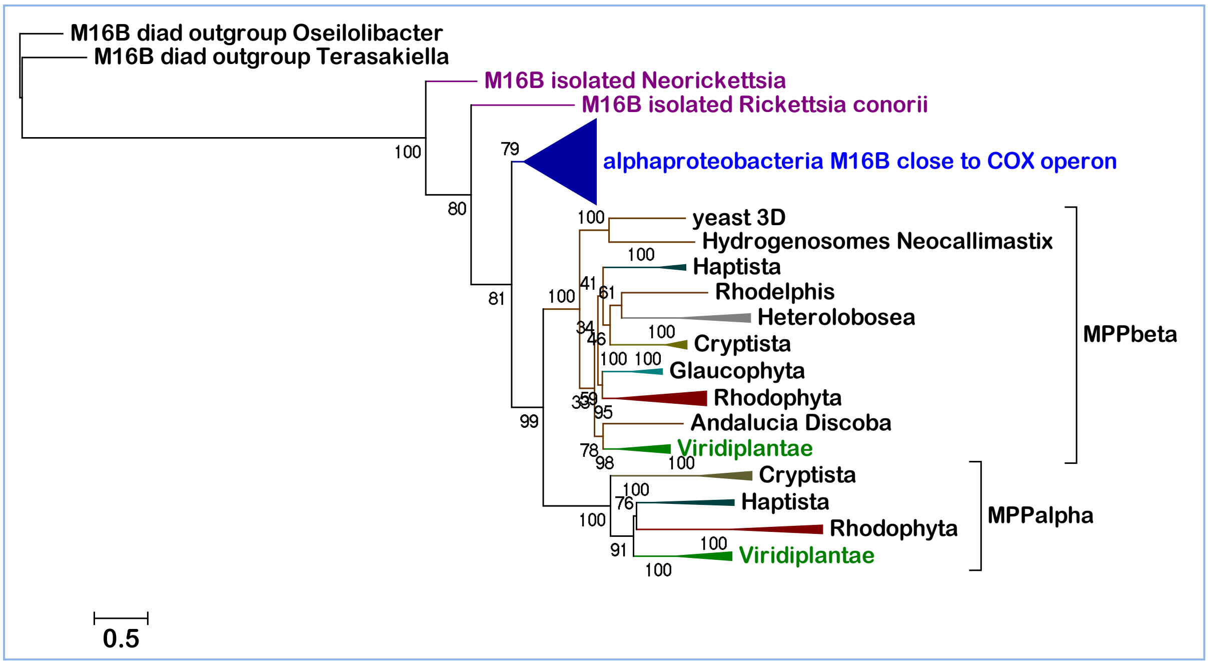

**Supplementary Figure S1**

**A.** Synteny COX11-COX3 in the mtDNA of early-branching Eukaryotes, with progressive distance due to the insertion of additional genes. **B.** Phylogenetic tree of M16B and MPP proteins. The ML tree was reconstructed from an alignment of equal number (33) of bacterial M16B and eukaryotic MPP proteins, using the best fit model LG obtained with the IQ-Tree web server system [43]. Note that the isolated M16B of *Rickettsia conorii* has been experimentally found to cleave mitochondrial pre-sequences as eukaryotic MPP proteins [21]. The outgroup proteins belong to the UPB group that forms part of a syntenic diad of M16B metallopeptidases, which are distinct from the M16.019 group closest to eukaryotic MPPbeta [22]. Similar trees were obtained with enlarged alignments and the EX_EHO mixture model.

#### Sneathiella sp. NORP273

| Accession | length(aa) | Protein description |
|---|---|---|
| MBL4740205.1 | 191 | YqgE/AlgH family protein |
| MBL4740206.1 | 194 | GNAT family N-acetyltransferase |
| MBL4740207.1 | 420 | **M16 peptidase** insulinase family protein |
| MBL4740208.1 | 466 | threonine synthase |
| MBL4740209.1 | 495 | carboxypeptidase M32 |
| MBL4740210.1 | 238 | **SURF1** family protein |
| MBL4740211.1 | 282 | oxygen-dependent coproporphyrinogen oxidase |
| MBL4740212.1 | 158 | tRNA (cytidine(34)-2\\'-O)-methyltransferase |
| MBL4740213.1 | 175 | ubiquinol-cytochrome c reductase iron-sulfur subunit **Rieske ISP** |
| MBL4740214.1 | 412 | **cytochrome b** N-terminal domain-containing protein |
| MBL4740215.1 | 255 | **cytochrome c1** |
| MBL4740216.1 | 291 | S-methyl-5\\'-thioadenosine phosphorylase |

**M16B** is five genes away from **ISP.** Note that the rest of the COX operon is missing, while a full subtype type a-III COX operon is present in the genome.

#### Alphaproteobacteria J084 MAG, Iodidimonadales

| Accession | length(aa) | Protein description |
|---|---|---|
| RMF12892.1 | 135 | flagellar basal body rod protein FlgC |
| RMF12879.1 | 197 | **cytochrome b**, partial |
| RMF12876.1 | 185 | ubiquinol-cytochrome c reductase iron-sulfur subunit **Rieske ISP** |
| RMF12877.1 | 405 | Sugar lactone lactonase YvrE |
| RMF12878.1 | 306 | M16B insulinase family protein, *partial* |
| RMF12860.1 | 356 | hypothetical protein D6757_08950 |

Partial **M16B** is one gene away from **ISP.**

#### Alpha proteobacteria Q-1 Iodidimonadales

| Accession | length(aa) | Protein description |
|---|---|---|
| GAK34097.1 | 272 | **cytochrome c1** |
| GAK34098.1 | 418 | **cytochrome b** |
| **GAK34099.1** | 181 | ubiquinol-cytochrome c reductase iron-sulfur subunit **ISP** |
| **GAK34100.1** | 436 | **M16B PqqL peptidase** (putative zinc protease) |
| GAK34101.1 | 248 | **SURF1** surfeit locus protein 1 |
| GAK34102.1 | 162 | DUF983 family protein with 2 closely spaced TM |
| GAK34103.1 | 268 | cytochrome c oxidase subunit 3 **COX3** |
| GAK34104.1 | 205 | cytochrome c oxidase assembly protein **CtaG_Cox11** |
| GAK34105.1 | 40 | **Cox4-like** hypothetical protein AQ1_02001 1TM |
| GAK34106.1 | 314 | protoheme IX farnesyltransferase **CtaB** |
| **GAK34107.1** | 544 | cytochrome c oxidase subunit **1 COX1** |
| GAK34108.1 | 308 | putative cytochrome c oxidase subunit 2 **COX2** |

**M16B** and **ISP** are adjacent as in *Iodididimonas muriae.*

**Precedent** in Eukaryotes is the filarial protein in Fig. 1: OZC11663.1 ubiuinol-cytochrome c reductase, iron-sulfur subunit of *Onchocerca flexuosa*

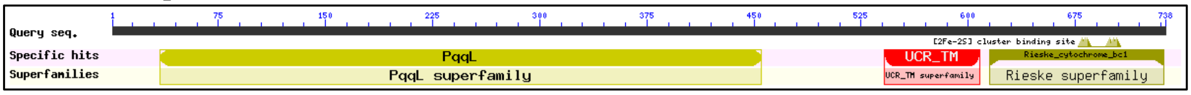

**M16B** **ISP**

**Supplementary Figure S2**. M16B-ISP genome contiguity.

#### A Lineage-specific distribution of anaerobic traits - OFORs

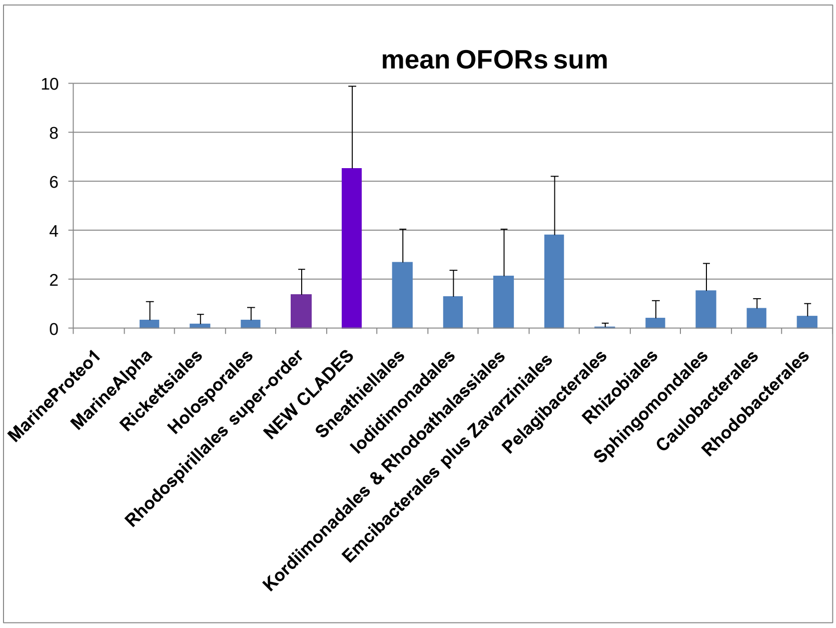

from aerobicplusSPY&kynU
5 May

Supplementary new Fig. 3AB 9 May

#### B comparative distribution of aerobic and anaerobic traits

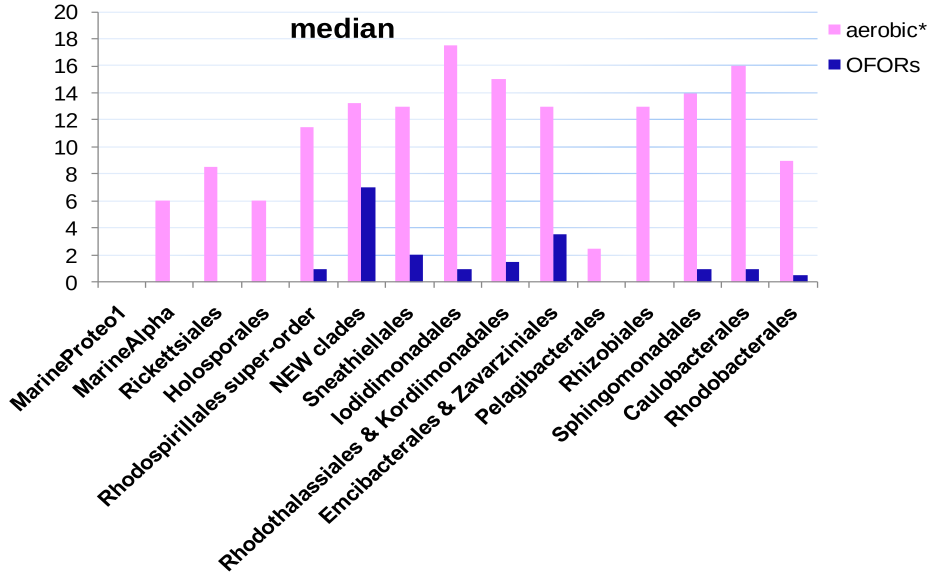

**Supplementary Fig. S3. Distribution of OFOR anaerobic traits.**
**A.** The lineage-specific distribution of the sums of all 2-oxoacid ferredoxin oxido reductases (OFORs) - not related to photosynthesis - is represented in mean plus SD values (cf. Figs. 5 and 6). **B.** Comparison of the distribution of cumulative scores for aerobic traits (cf. Fig. 2) and the sums of OFORs anaerobic traits along the lineages of alphaproteobacteria (Supplementary Table S2, rightmost column).*****The median values were subtracted by 6, the median of aerobic scores for gammaproteobacteria (Fig. 2).

#### A Major indels in COX3 proteins

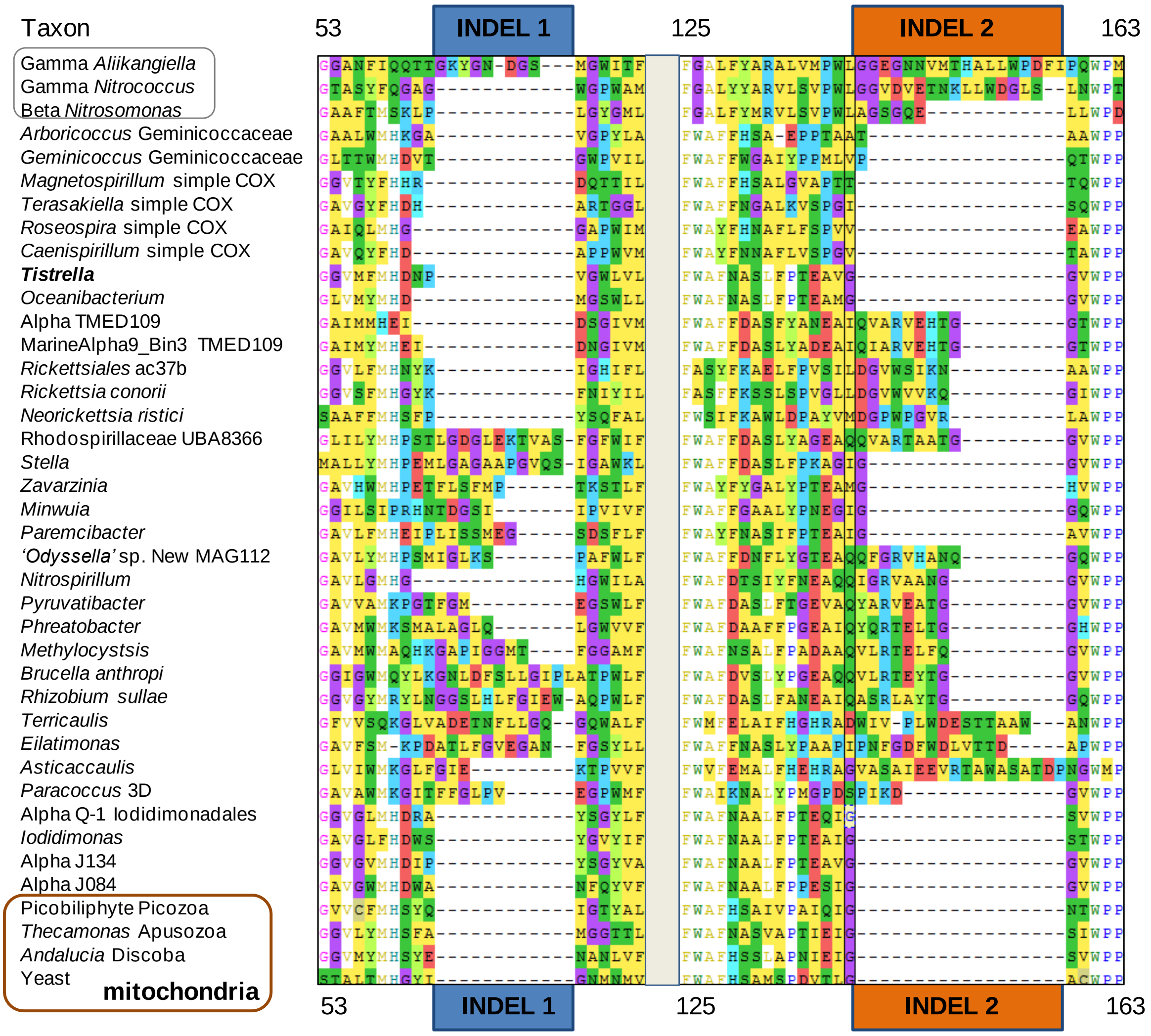

**Supplementary Fig. S4A**. **Major INDELs in COX3 sequences**. The number before and after the INDELs correspond to the position in the alignment of diverse bacterial (top) and mitochodnral (bottom) COX3 proteins.

### B Mapping conserved INDELs of COX3 along a phylogenetic tree

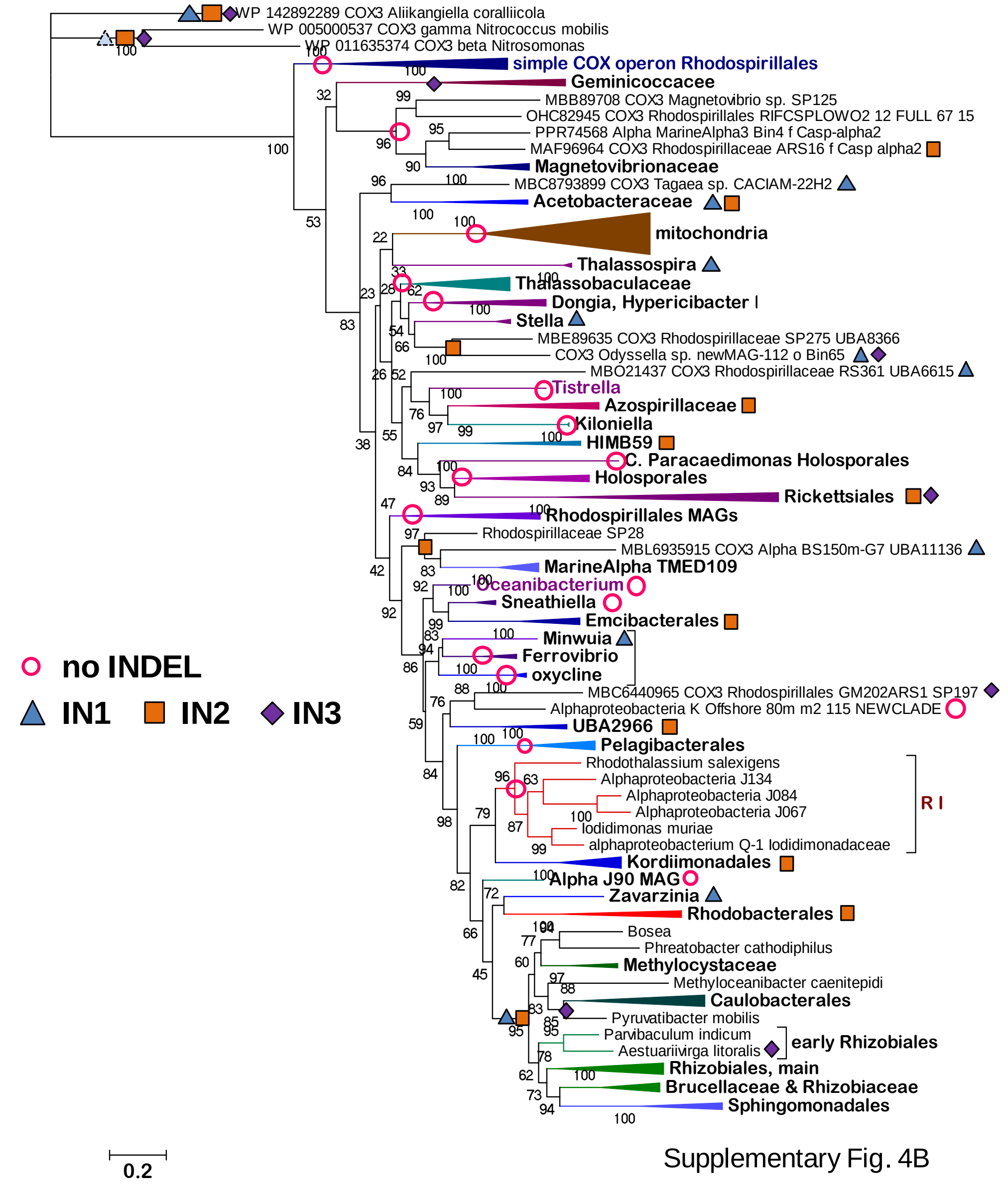

**Supplementary Fig. S4B**. Mapping of the INDELs of COX3 along a ML tree of the protein.
The ML tree was reconstructed from an alignment of 140 COX3 sequences equivalent to that used in part A. It is representative of 20 similar trees obtained with different models and slighlty different combinations of aligned proteins.

**Supplementary Table S1.** MPP and related proteins in various eukaryotic taxa.

Original file: supplementaryMPPCPlist.xls .

| **taxon** | **classification** | **beta-MPP** | **CP-1** | **alpha-MPP** | CP-2 | **other** |
|---|---|---|---|---|---|---|
| ***Cyanidioschyzon merolae*** | Archaeplastida, Cyanidiaceae | XP_005537012 & XP_005536303 different | | XP_005536642 | | XP_005536303 |
| ***Cyanidiococcus yangmingshanensis*** | Archaeplastida, Cyanidiaceae | KAF6002968 | | KAF6002367 partial | | KAF6002180 & KAF6002181 part. |
| ***Gracilariopsis chorda*** | Archaeplastida, Florideophyceae | PXF48706 | | PXF49250 | | |
| ***Chondrus crispus*** | Archaeplastida, Florideophyceae | XP_005715652 | | XP_005717973 | | |
| ***Porphyra umbilicalis*** | Archaeplastida, Bangiophyceae | OSX72742 & OSX76050 different | | OSX81301 | | |
| ***Porphyridium purpureum*** | Archaeplastida, Bangiophyceae | KAA8497343 | | | KAA8499490 likely by **LGT** | 5 doublet & KAA8500059 |
| ***Compsopogon caeruleus*** | Archaeplastida, red algae, Compsopogonophyceae | CAD9221014 & CAD9221016 | | CAD9237020 | | |
| ***Galdieria sulphuraria strain 074W*** | Archaeplastida, red algae, Cyanidiaceae | XP_005703305 | | XP_005709111 | | XP_005708473 |
| ***Rhodelphis limneticus*** | Rhodelphida | **g5912** | | **g989** | | |
| ***Gloeochaete wittrockiana*** | Archaeplastida, Glaucophyta | CAE5718633 | | | | |
| ***Cyanoptyche gloeocystis*** | Archaeplastida, Glaucophyta | CAD9553425 | | | | |
| ***Trebouxia sp. A1-2*** | Archaeplastida, Chlorophyta | KAA6428238 | | KAA6416852 partial | | |
| ***Ostreococcus tauri*** | Archaeplastida, green algae, Mamiellophyceae | XP_022841363 & OUS46027 | | XP_003084396 | | 5 doublets |
| ***Prasinoderma coloniale*** | Viridiplantae, Prasinodermophyta | CAD8249806 partial | | CAD8249804 | | at least 1 doublet |
| ***Chlamydomonas reinhardtii*** | Archaeplastida, Chlorophyta, Chlamydomonadales | XP_001697021 | | XP_001697130 | XP_001691043 | 3 doublets |
| ***Klebsormidium nitens*** | Archaeplastida, Streptophyta, Klebsormidiophyceae | GAQ80207 | | GAQ89643 | | 1 M16 long GAQ91500 |
| ***Chara braunii*** | Archaeplastida, Streptophyta, Charophyceae | GBG76521 | | GBG84704 | | 5 doublets & GBG89 |
| ***Wollemia nobilis*** | Archaeplastida, Streptophyta, Pinopsida | JAG87885 | | JAG87957 | | 2 doublet M16 |
| ***Vigna radiata*** | Archaeplastida, Streptophyta, Fabaceae | **XP_014498085 X1*** XP_014516218 & XP_014498086 X2 | | **XP_014516769*** XP_014505465 & XP_022636067 | | 4 doublets |
| ***Guillardia theta CCMP2712*** | Cryptophyceae, Pyrenomonadales | XP_005841328 | CAE2270003 partial | XP_005835389 | CAE2321217 | |
| ***Rhodomonas abbreviata*** | Cryptophyceae, Pyrenomonadales | CAE1998097 | CAE1991465 partial | CAE1988489 | CAE1961634 | |
| ***Cryptomonas curvata*** | Cryptophyceae, Cryptomonadales | CAD8644786 & CAD8649381 partial | CAD8630992 long | CAD8656898 | | |
| ***Palpitomonas bilix*** | Eukaryota incertae sedis | CAE0259888 partial | | | | CAE0259888 Ptr fragment |
| ***Andalucia godoyi*** | Discoba, Jakobida | KAF0852499 | | KAF0852193 | | |
| ***Percolomonas cosmopolitus*** | Discoba, Heterolobosea | CAD9077212 | | CAE2765079 | | |
| ***Naegleria gruberi*** | Discoba, Heterolobosea | EFC43189 or XP_002675933 | XP_002673612 or beta isoform? | XP_002669264 | | |
| ***Naegleria fowleri*** | Discoba,Heterolobosea | KAF0975433 | | KAF0975495 | | |
| ***Bigelowiella natans*** | SAR, Rhizaria, Chlorarachniophyceae | CAE3743251 | | CAE3739956 & CAE2012412 | | |
| ***Spongospora subterranea*** | SAR, Rhizaria, Endomyxa | CRZ09285 | | CRZ09126 | | |
| ***Chrysochromulina tobinii*** | Haptophyta | KOO21062 | | KOO21553 | | |
| ***Emiliania huxleyi*** | Haptophyta | XP_005762229 XP_005790187 | several possible | CAG1280390 | several possible | |
| ***Pavlova gyrans*** | Haptophyta | CAE4048132 | CAE4043026 partial | CAE4038285 | | |
| **taxon** | **classification and super-group** | **beta-MPP** | **CP-1** | **alpha-MPP** | **CP-2** | **other** |
| * in 3D structure of complex III | | | | | | |
